# Fibroblast Density is a Risk Factor for Drug-induced Arrhythmias

**DOI:** 10.1101/2025.01.19.633080

**Authors:** Kayo Hirose, Shinjiro Umezu, Daisuke Sato

## Abstract

A recent study by Kawatou *et al*. has shown that the local heterogeneity of ion channel conductance is a critical substrate for focal or reentrant arrhythmias. However, the role of fibroblasts with repolarization heterogeneity in the initiation and maintenance of arrhythmias remains unknown. In this study, we investigated how diffuse fibrosis contributes to the formation of focal and reentrant arrhythmias under drug-induced heterogeneity using physiologically detailed mathematical models of the human heart. To simulate drug-induced heterogeneity, we varied the maximum conductance of transmembrane potassium and calcium currents, leading to heterogeneity in action potential duration (APD). Then, we assessed the effects of different fibrosis densities (FD) on the occurrence of premature ventricular complexes (PVCs). Fibroblasts were randomly and evenly inserted into the tissue, and various FD levels ranging from 0 to 35% were examined. We found a biphasic relationship between FD and drug-induced PVCs. Within a certain range of FD, FD positively correlated with PVC susceptibility. However, excessively high fibrosis levels were associated with reduced susceptibility to PVCs. In addition, the self-sustainability of arrhythmias exhibited a positive correlation with FD. This study demonstrates the interplay between the diffuse fibrosis and the drug-induced heterogeneity of APD in the genesis of ventricular arrhythmias.

**Author summary:** Sudden cardiac death remains a leading cause of death worldwide. Understanding the mechanisms underlying arrhythmia and its precursors is critical for the development of effective therapies and drugs. Repolarization heterogeneity plays a crucial role in both the initiation and maintenance of arrhythmias. Fibroblasts constitute a vital component of cardiac structure, originating from the remodeling of ventricular wall cells or the transformation of injured myocardial cells. Fibroblasts are known to couple with and alter the electrical properties of myocardial cells. However, our understanding of the role of fibroblasts in the development of arrhythmia remains limited. In this study, we employed a physiologically detailed mathematical model of cardiac tissue to investigate the roles of drug-induced heterogeneity and diffuse fibrosis in the initiation and maintenance of arrhythmias. We used 2D and 3D computational models to simulate various levels of drug-induced heterogeneity conditions with normal to pathological levels of fibroblast density (FD). We found that within a certain range of FD, fibroblasts promote PVCs under drug-induced heterogeneity. However, if FD exceeds 30%, the occurrence of PVCs decreases (biphasic relationship). On the other hand, the self-sustainability of VF (ventricular fibrillation) consistently increases with FD. This study implies that fibroblasts in cardiac tissue may play different roles in the initiation and maintenance of arrhythmia.

## Introduction

Sudden cardiac death (SCD) is a leading cause of death in many countries. In the United States alone, SCD accounts for approximately 184,000 to 400,000 deaths every year [1]. Ventricular fibrillation (VF) induced by polymorphic ventricular tachycardia (PVT) is a common mechanism of cardiac arrest that causes SCD. PVT presents as a rapid and regular ectopic rhythm, while VF manifests as a chaotic and irregular quivering rhythm. PVT can evolve from ventricular tachycardia under certain conditions. Currently, widely accepted explanations for the mechanism of ventricular fibrillation include the multiple wavelet hypothesis [2], high-frequency focal source hypothesis [3], and spiral wave break-up hypothesis [4]. However, due to the diverse manifestations of ventricular fibrillation, which may be influenced by different mechanisms, there is currently no hypothesis that can explain all observed phenomena [1]. What is currently known is that changes in cardiac repolarization reserve (RR) and heterogeneity in effective refractory period are crucial for the initiation and maintenance of VF [5]. RR represents the ability of the tissue or myocyte to resist interference to achieve orderly and rapid repolarization, and a reduction in outward current (e.g., I_Kr_ or I_Ks_) and/or an increase in inward current (e.g., I_CaL_) of myocytes can lead to the reduction of RR [6]. The reduced RR leads to prolonged action potential duration (APD) and increases the propensity to trigger early afterdepolarization (EAD), the secondary voltage depolarizations during the repolarizing phase of the AP. EAD is considered closely related to the occurrence of many kinds of ventricular arrhythmia, such as PVT, VF and TdP [7][8][9][10][11]. The heterogeneity in APD (and thus effective refractory period) is primarily attributed to the spatial structural heterogeneity of cardiac tissue. But more importantly, it changes dynamically due to various factors such as pacing rates, posttranslational modifications, and drug administrations. For example, compared to epicardial and endocardial cells, M cells are more sensitive to a variety of pharmacological agents that can change I_Kr_, I_Ks_, and I_CaL_ [12]. This sensitivity is reflected in the greater APD prolongation of M cells than endocardial and epicardial cells following the blockade of I_Kr_, I_Ks_, and augmentation of I_CaL_. The heterogeneity of APDs can provide a substrate for ectopic beats and reversal of ventricular wall activation direction, thereby leading to the occurrence of VT/VF [13].

Fibroblasts play a crucial role in maintaining the normal structure and function of the heart, and they actively participate in the remodeling of the ventricular wall under pathological conditions, such as myocardial infarction and pressure overload. In fact, myocardial fibrosis accompanies most cardiac pathological conditions [14]. Although in most myocardial diseases, the extent of cardiac fibrosis predicts adverse outcomes, fibrosis is not necessarily the primary cause of dysfunction. The adult mammalian heart has negligible regenerative capacity and heals through the formation of a scar. Thus, in many cases, cardiac fibrosis is reparative, reflecting the replacement of dead cardiomyocytes with a collagen-based scar. The fibrosis thus formed impairs the propagation of electrical impulses and generates reentrant circuits, which may contribute to arrhythmias. In fact, the association between fibrosis and increased risk of ventricular arrhythmias has been demonstrated in virtually every cardiac condition for which fibrosis remodeling has been evaluated.

Fibroblasts can electrotonically couple with myocytes and contribute to the electrical properties of the myocardium. In vitro and in silico studies have shown the effects of fibrosis on myocardial electrical function [15]. Modeling studies have shown that APD of a single myocyte is shortened, and EAD activity is impeded when coupling to fibroblast [16][17]. However, experiments demonstrate that the occurrence of EAD and ectopic triggers more frequently originated in areas with relatively higher FD in aged rats and rabbits [18], and a recent experiment shows coupling of fibroblasts and myocytes may be responsible for cardiac arrhythmias [19]. The gap junction coupling between fibroblasts and myocytes can help or hinder the PVC triggering ability of EAD masses [20]. These EADs have been known to trigger arrhythmia, but clear understanding of the influence of fibrosis on VF remains lacking and it is still unclear what role the fibrosis plays in the initiation and maintenance of VT/VF.

According to the previous study, the coupling of fibroblasts and cardiomyocytes can be seen as the source and the sink in the electric field [21]. When the myocytes are undergoing the depolarization period, the exciting myocytes can be regarded as the current source, at the same time, the sink, fibroblasts which connected to the myocytes will divert part of the current, lead to the decrease of excitement potential. As for the repolarization period, the situation becomes opposite. The effect of fibrosis on PVC triggering also reported [22][23]. It shows a promoting effect at 10% FD and an inhibiting effect at 15% FD. They attribute the effect of fibrosis on PVC triggering to a source-sink mismatch.

However, in the presence of heterogeneous regions, fibrosis has distinct effects on M cells, Endo, Epi, and heterogeneous region M cells, resulting in complex mechanisms for the generation and maintenance of ectopic beats and re-entrant waves. This study investigates how drug-induced heterogeneous substrates at different fibrosis levels lead to PVC and sustained VF.

## Materials and methods

To investigate the effects of fibroblasts on the initiation and maintenance of VF/VT, we used mathematical models. We used 2D tissue models to investigate the underlying mechanisms under normal to pathological levels of RR and fibrosis distributions. We also use an anatomical ventricle model to verify the validity of the mechanisms in 3D space.

### Cardiac action potential model

In this study, we used a physiologically detailed model of the human cardiac action potential model (Tusscher-Noble-Noble-Panfilov (TNNP) 2006) [24]. The membrane potential of the myocyte (V_m_) is governed by

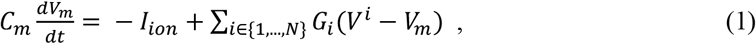

where C_m_ is the membrane capacitance, which has a value of 185 pF. *N* indicates the number of neighbor cells around the myocyte. *V*^*i*^ refers to the membrane potential of the i^th^ neighbor cell. *G*_*i*_ is the gap junction conductance between the i^th^ cell and myocyte. *I*_*ion*_ is the sum of all transmembrane ionic currents of the myocyte:

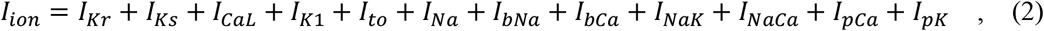

where *I*_*Kr*_ is rapid delayed rectifier current, I_Ks_ is slow delayed rectifier current, *I*_*CaL*_ is L-type Ca^2+^ current, *I*_*K1*_ is inward rectifier K^+^ current, I_to_ is transient outward current, *I*_*Na*_ is fast Na^+^ current, *I*_*bNa*_ *I*_*NaCa*_ is Na^+^/Ca^2+^ exchanger current, *I*_*bCa*_ and *I*_*bNa*_ are background Ca^2+^ and Na^+^ currents, *I*_*NaK*_ is Na^+^/K^+^ pump current, *I*_*pCa*_ and *I*_*pK*_ are plateau Ca^2+^ and K^+^ currents, respectively.

Fibroblasts are modeled based on MacCannell *et al* [16]. The gap-junctional conductance between the fibroblast and myocyte *G*_gap_ was set to 3 nS. The membrane capacitance *C*_*f*_ of fibroblasts is measured in the range of 6-10 pF [25][26]. The experimentally measured resting potential *E*_*f*_ is between -10 and -50 mV [27]. In this study, we use the same parameters as MacCannell, *et al* [16]. The *C*_*f*_ was set to 6.3 pF, and the *E*_*f*_ was set to -50 mV. The membrane conductance G_f_ was set to 0.2 nS. The membrane potential V_f_ of the fibroblast is governed by

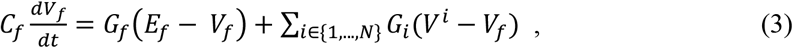

where *N* is the number of neighbor cells around the fibroblast. *V*^*i*^ is the membrane potential of the i^th^ neighbor cell. *G*_*i*_ is the gap junction conductance between the i^th^ cell and fibroblast.

### Cardiac tissue models

To investigate the mechanisms of initiation and maintenance of VF and its relationship with cardiac fibrosis, we employed both 2D and 3D models in this study. The 2D model was utilized to explore the fundamental mechanisms underlying PVC formation. Its simplified structure allowed for a detailed examination of the electrophysiological properties and regulatory mechanisms associated with VF. In contrast, the 3D model was employed to simulate the study in a more realistic cardiac environment. By considering the three-dimensional structure of the heart and incorporating the presence of fibrosis, the 3D model better replicated actual conditions, enabling a comprehensive analysis of the impact of fibrosis on VF. This combined approach of utilizing both 2D and 3D simulations aimed to provide a more integrative understanding of VF mechanisms and the effects of fibrosis, contributing valuable insights into clinical implications and interventions.

### 2D tissue model

The 2D tissue model represents a transmural section of the ventricle, and its geometry is shown in Fig 1 and Fig 2. The tissue was modelled as a rectangle consisting of 512×100 nodes (physical size of 12.8cm × 2.5cm). The tissue incorporates, three cell types, namely, epicardial (epi), midmyocardial (mid), and endocardial (endo) cells. The ratio of the thicknesses of the endo, mid, and epi cell areas is set at 0.3:0.45:0.25. To simulate diseased conditions, we reduced repolarization reserve by decreasing the outward currents (*I*_*Kr*_ and *I*_*Ks*_) and increasing the inward current (*I*_*CaL*_). Considering the impact of heterogeneity on PVCs, we changed two areas’ conductance of *I*_*CaL*_ in the myocyte part of tissue model, in these areas, the conductance of *I*_*CaL*_ channel is set higher than other cells, and the repolarization reserve in this area was significantly reduced. In the heterogeneous area, affected by reduced *I*_*Ks*_ and increased *I*_*CaL*_, the APD of myocyte cells in this area will be significantly longer than the surroundings. This creates spatial and temporal differences in cell repolarization that can help trigger PVCs and prolong the arrhythmic state. Details are shown in the supporting materials.

**Fig 1.**
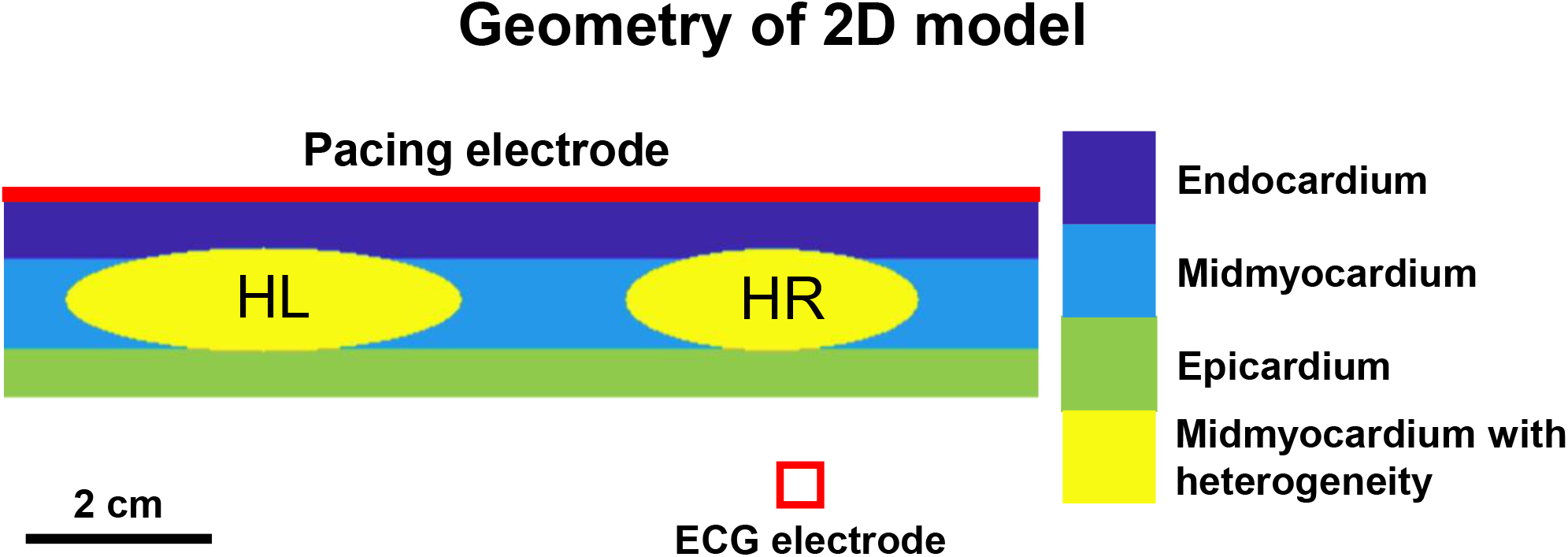
Geometry of 2D model. the 2D tissue model is composed of 4 kinds of cells: Endocardium (Endo), Midmyocardium (Myo), Epicardium (Epi) and Midmyocardium with heterogeneity. The cell area of Endo, Myo and Epi is 0.3:0.45:0.25. In the model, the electrodes were placed at 2.5 cm above the surface of the Epi and the surface of Endocardium. The conductance of I_CaL_ definitized as different value in the tissue of Midmyocardium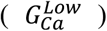 and Midmyocardium with Heterogeneity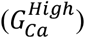, and 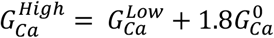.

**Fig 2.**
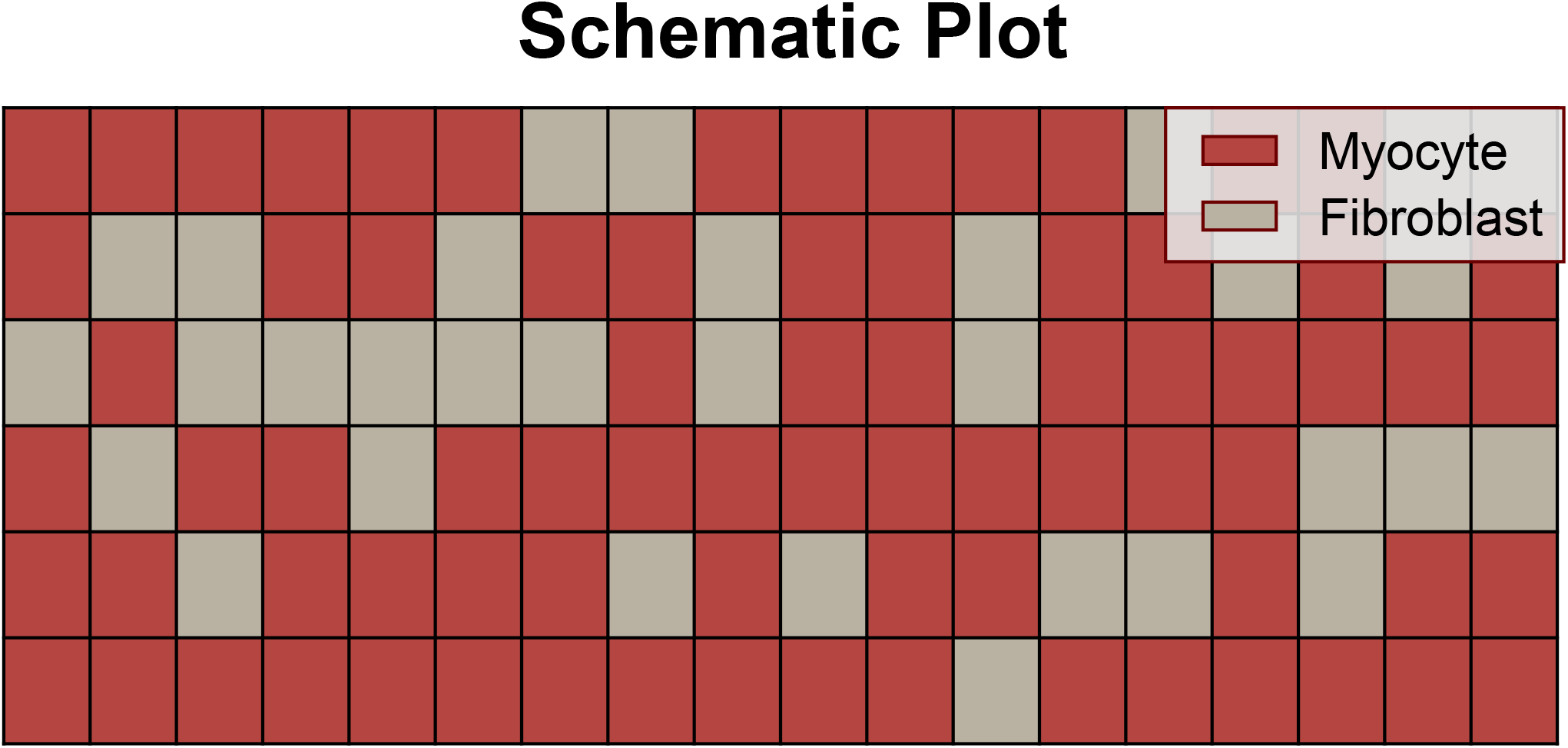
Schematic Plot The 2D model. The whole model is consisting of 512×100 grids, the figure shows part of the model. We assume that cardiac cells and fibroblasts are the same size, and that fibroblasts are randomly distributed among all cardiac cells. The figure shows the state of diffuse fibrosis (FD) = 30%.

In all 2D simulations, the spatial and temporal resolutions were set to be δx = 0.25 mm and δt = 0.02 ms, respectively. The myocytes are coupled diffusively with a diffusion coefficient of *D*_mm_ = 0.00154 cm^2^ms^-1^. The conduction velocity (CV) in tissue without fibroblasts was 68 cm/s. The coupling strength between fibroblasts was set to one-tenth of *D*_mm_ [28]. The stimulus was applied from the pacing electrode at a frequency of 1.4 Hz.

### 3D human ventricular model

To build the 3D human ventricular model, we first resampled the 0.35mm resolution mesh of a publicly available human cardiac model (mesh 01) [29] to an average edge length of 2.06±0.8mm tetrahedral mesh using meshtool (https://bitbucket.org/aneic/meshtool/src/master/). The label and fiber orientation of the original model were retained in the resampled model. We extracted the ventricular part, which consists of 49,262 nodes and 231,937 elements. Then, we generated the Purkinje network using publicly available tools (https://github.com/fsahli/fractal-tree) developed by Sahli Costabal *et al*. [30]. The network consists of 7,876 line elements and 7,877 nodes. Similar to the 2D model, epi, mid, and endo cells were implemented. The ratio of the endo, mid, and epi cell area was set as 0.3:0.45:0.25, referring to the transmural label inherited from the original model. As shown in fig.2(d), we inserted one mid tissue with heterogeneity in the left ventricular (LV) and right ventricular (RV), respectively. We used the same settings for the conductance of the *I*_*Ks*_, *I*_*Kr*_, *I*_*CaL*_ currents as of 2D model. The fibroblast is randomly and evenly distributed throughout the entire tissue, and we varied FD from 0% to 35%.

In the 3D simulation, we used the time adaptive algorithm [31] with the time step δt varying between 0.01 ms and 0.1 ms. The longitudinal and transversal conductivities of myocytes were set As 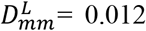 cm^2^ms^-1^ and 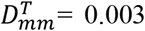 cm^2^ms^-1^. When there is no heterogeneity area and fibroblast in the ventricle, the approximate longitudinal CV is 0.74 m/s and transversal CV is 0.26 m/s which are consistent with the experimental measurements [32]. Similar to the 2D simulation, the coupling strength between fibroblasts was set to one-tenth of 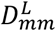 and 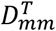. Purkinje fibers were modeled as a bifurcated one-dimensional cable model embedded in the mesh of a 3D ventricular model, with the fixed conduction velocity set at 3 m/s [33]. The pacing stimulus is released from the Purkinje fiber, and its frequency is 1.4 Hz. The activation time map of the Purkinje network is shown in the Supplemental figure 5.

### Numerical calculation, implement and Pseudo-ECG computation

Both 2D and 3D models were solved by the forward Euler integration scheme. The numerical solver was implemented with MATLAB programming languages. Computations were performed with double precision and ran on an Intel Core Xeon(R) Silver 4214 CPU with NVIDIA Quadro P620 graphics cards.

In the 2D simulation, the pseudo-ECGs were calculated by the formula:

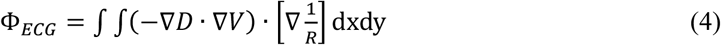

Where D is the diffusion tensor, V is the voltage, and R is the distance from each point of the tissue to the ECG electrode. The ECG electrode was placed 2.5 cm from the epicardial side surface of the tissue.

In the 3D simulation, the pseudo-ECGs were calculated using the similar formula:

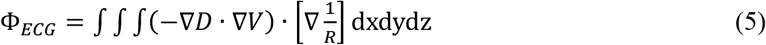

R is the distance from each point of the ventricular to the ECG lead. The ECGs of 3D were computed with leadIwhich is shown in Fig 3C. The fibroblasts were not taken into ECG calculation in both 2D and 3D simulations.

**Fig 3.**
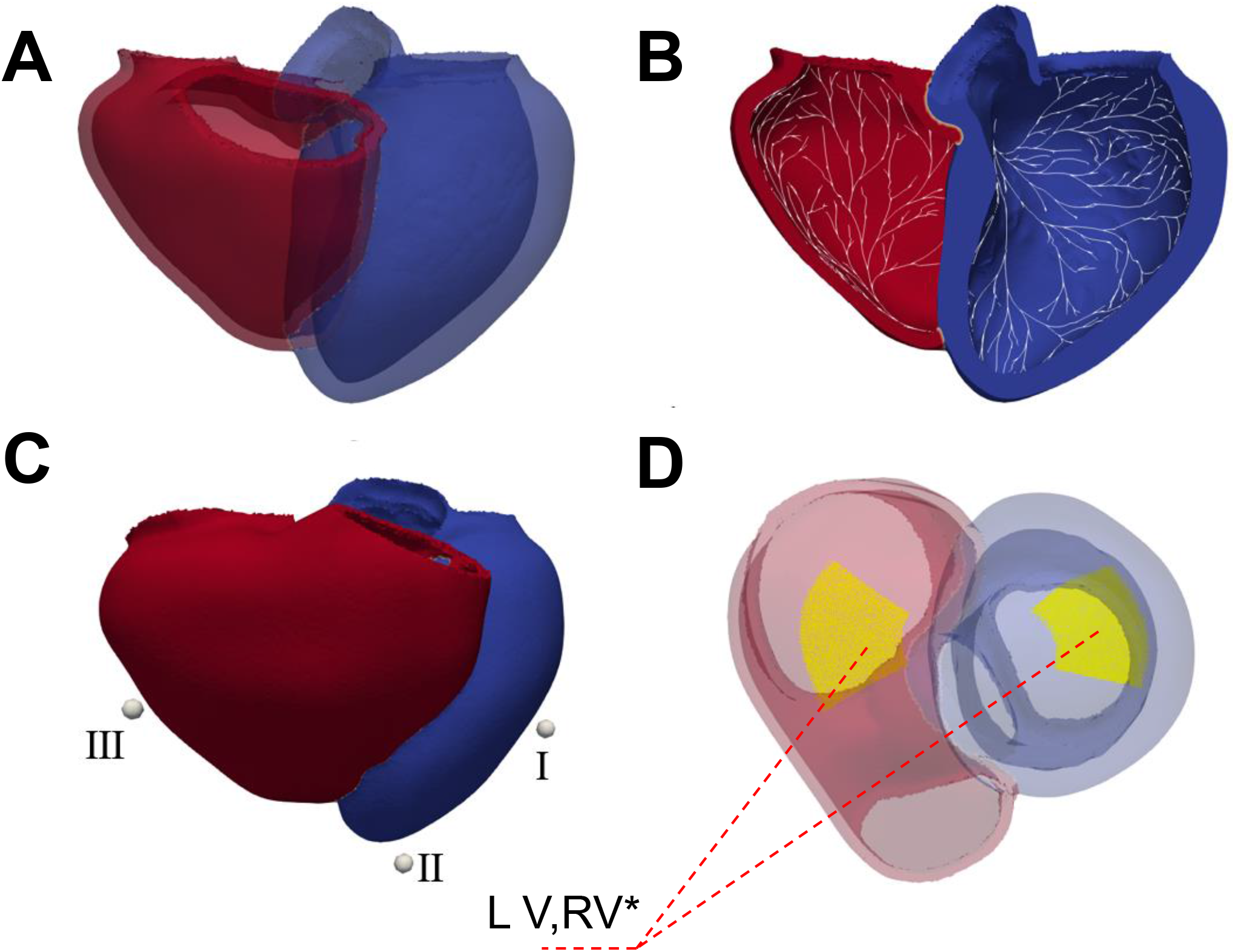
Geometry of 3D model. (a) human ventricular, the blue chamber is left ventricular, the red chamber is right ventricular. (b) ventricular with Purkinje network, the while line shows the Purkinje network. (c) lead placement for pseudo-ECG, the white spheres are electrode of leadI, leadII, lead III. (d) The yellow bulks indicate there are heterogeneous areas where the conductivity of L-type calcium channels is higher than that of the surrounding tissue to trigger PVC and maintain VF.

## Results

### APD was prolonged as FD increased

Since prolonged APD is a significant manifestation of EADs, which are associated with PVCs and arrhythmias [19], we first investigated how fibrosis affects APD in tissue. We measured changes in APD in heterogeneous areas in Fig 1 under various FD (Fig 4), ranging from 0% to 30%. We found that the APD was prolonged as FD was increased. With the increase of FD, both median and mean values of APD_90_ are increased, indicating greater propensity for EADs. In addition, variability in APD was also increased as FD was increased, indicating heterogeneity in APD can be increased in this area. It is worth noting that when FD exceeded 30%, the maximum APD in tissue extended over 700ms. As a result, some myocytes still not fully repolarized when the next depolarization wave comes at a pacing frequency of 1.4 Hz (∼cycle length≈714ms). In these cases, the repolarization of these cells may be interrupted by the next waves, and electrical stimulation caused by the excitement of surrounding cells can reopen *I*_*CaL*_ channels, leading to EADs or DADs (delayed afterdepolarizations) as shown in Fig 4 and Fig 8. These myocytes cause slow conduction or block during the subsequent pacing. Moreover, the increasing APD_90_ - APD_30_, which shows the extent of action potential triangulation [8], indicates that the time window over which tissue could be re-excited in final repolarization is widened. This property contributes to the propensity for EADs [34] that can induce the ectopic beats that initiate and maintain VF.

**Fig 4.**
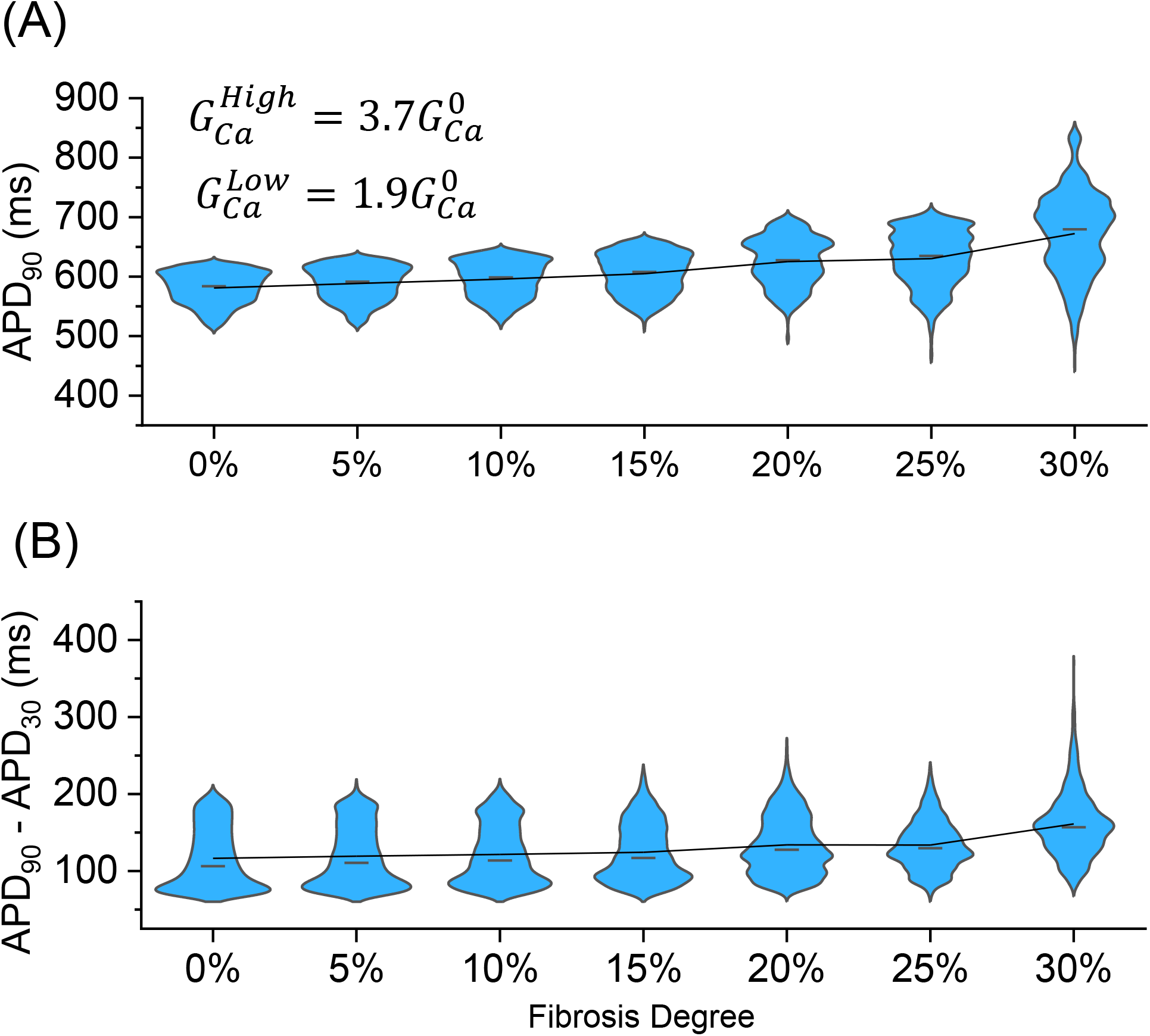
APD_90_ *^1^(a) and APD_90_ -APD_30_ *^2^(b) of myocytes in mid tissue with heterogeneity. The black connection line is the mean value, the black horizontal line inside the box is the median value. The simulation time is 800ms.

**Fig 5.**
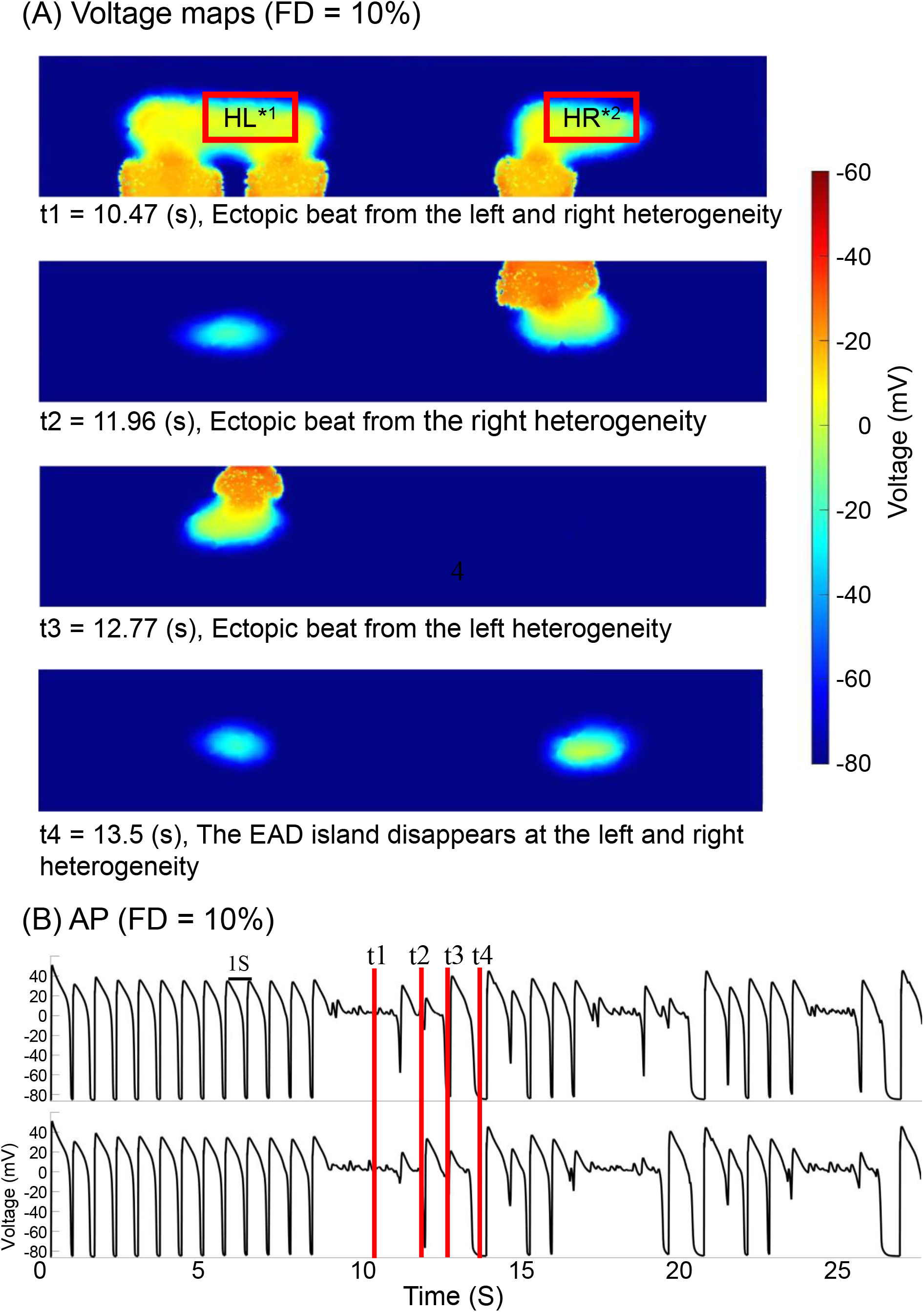
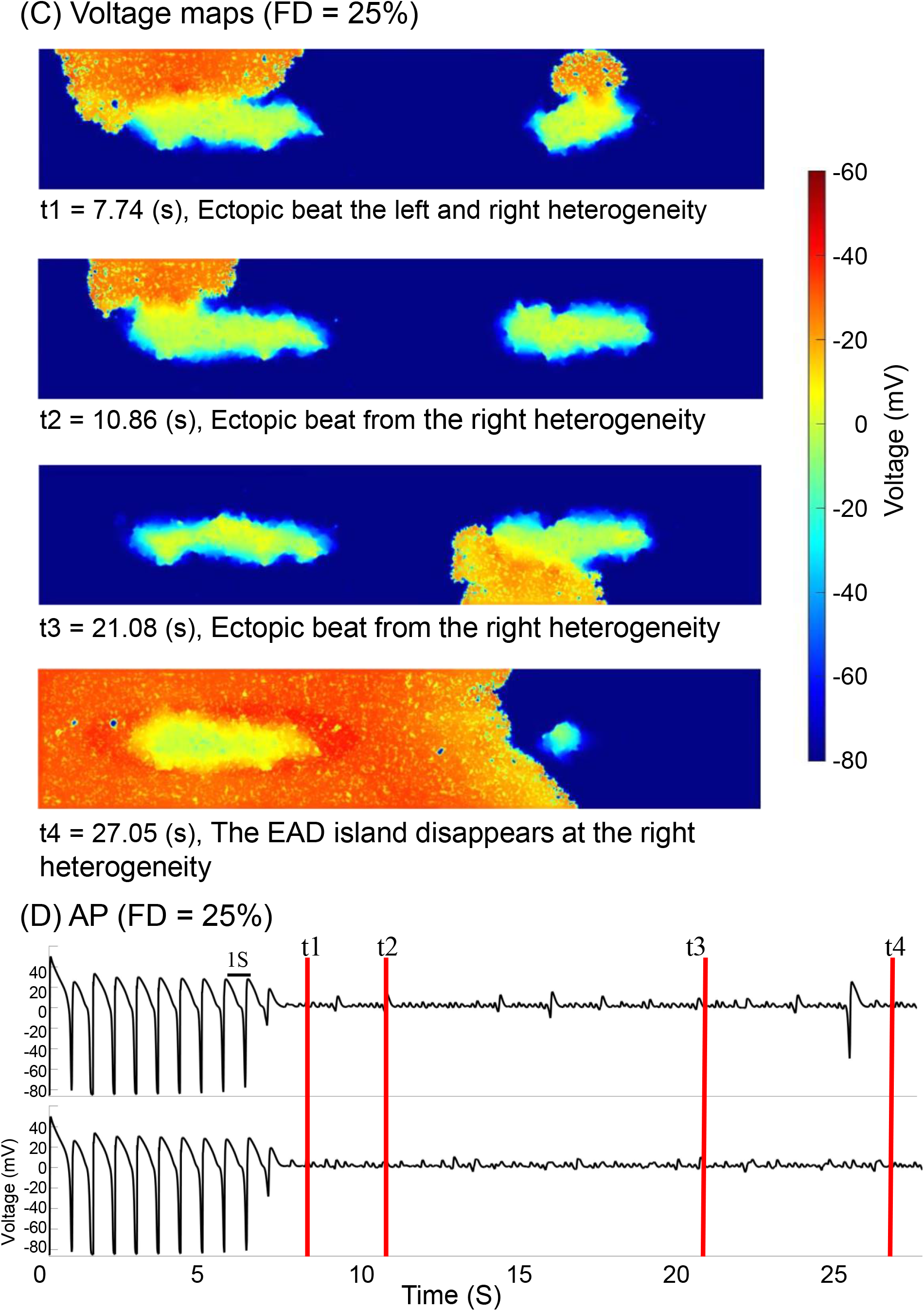
Voltage maps and AP of 2D model in 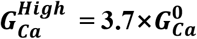 and 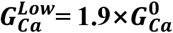, FD = 10% and FD =25%. (A) Voltage maps (FD = 10%). HL and HR represent the heterogeneous regions on the left and right sides of the tissue model, respectively. (B) AP of monitor point at left and right heterogeneity (FD = 10%). (C) Voltage maps (FD = 25%). (D) AP of monitor point left and right heterogeneity (FD = 25%)

### Fibroblasts promote EAD formation

The simulations above have shown that fibrosis can promote EAD formation in tissue. To understand how fibrosis affects EAD formation, we simulated a single cell coupled with fibroblasts.

The number of coupled fibroblasts ranges from 0 to 4, as shown in Fig 13. The same as the 2D tissue simulations, the parameters of the action potential model are from TNNP06. We also used the same drug-induced arrhythmia conditions: block the *I*_*KS*_, 25% block the *I*_*Kr*_ to attenuate the repolarizing current, thus reducing the RR and make the cells susceptible to EAD. We also reduced the RR by increasing the conductance of L-type calcium channels (G_Ca_) to increase the outward current (*I*_*CaL*_) during the plateau phase of the action potential. We call the cell with a small increase in *I*_*CaL*_ as the high RR condition 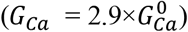(Fig 9A), and the cell with a large increase in the *I*_*CaL*_ as the low RR condition 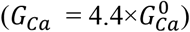(Fig 9B).

When RR is low and the action potential already has a long plateau, fibroblasts hardly affected the autonomous persistence of EAD, and only slightly affected the maximum potential and plateau potential (Fig 9B).

However, when RR is high, coupled with more fibroblasts, the effect on myocyte APD was biphasic (Fig 9A). In the absence of coupling with fibroblasts, we observed one oscillation (i.e EADs) in the plateau and returns to the resting potential. When one or two fibroblasts are added, the myocytes act as current sources and fibroblasts act as current sinks initially. However, in the late plateau phase, fibroblasts act as current source and prolongs the action potential and promotes EAD formation. When three or more fibroblasts are added, fibroblasts act as current sinks primarily, and reduce the membrane potential. Thus, we observed fewer EADs. Here, under high RR conditions, a small number of fibroblasts (i.e. low density in tissue) can promote EAD formation, while a large number of fibroblasts (i.e. high density in tissue) may facilitate the termination of EADs.

### Initiation and maintenance of VT/VF in the 2D model

We then investigated the effects of FD on the initiation and maintenance of VT/VF. Here we show the comparison between FD=10% (Figs. 5A and B) and FD=25% (Figs. 5C and D).

When FD is 10%, the first ectopic beat originates from HL (heterogeneous area on the left side of the tissue, see Fig 5) at 9.07 s. Subsequently, the following ectopic beat was generated at t = 9.79 s from the HL. At t = 9.81 s, another ectopic beat arose from the HR (heterogeneous area on the right side of the tissue, see Fig 5). Then at t = 10.47 s, two ectopic beats were created almost simultaneously from both HL and HR (Fig 5A). After two ectopic beats from HR (t = 11.96 s, Fig 5A) and HL (t = 12.77 s, Fig 5A) respectively, the PVCs terminated as EAD islands disappeared almost simultaneously (t = 13.5 s, Fig 5A). The tissue then returned to the normal pacing cycle and propagation. In this case, the action potential waves originating from the HL stimulated the HR and maintained the EAD island in the HR, causing subsequent ectopic beats. Conversely, the action potential wave from the HR due to this ectopic beat (t = 11.96 s fig 5A) stimulated the HL and maintained the EAD island in the HL, leading to further ectopic beats. This ping-pong like patterns can occur repeatedly. Only when both HL and HR fail to generate ectopic beats before the EAD islands disappear, the VT/VF like activity can cease spontaneously. This is a typical scenario when FD is moderate. A movie of this simulation with voltage maps along with ECG is provided as Movie S1.

When FD is high (FD 25%), ectopic beats arise simultaneously from both HL and HR (t = 7.74, as illustrated in Fig 5C). Subsequent ectopic beats emanate alternately from HL (t = 10.86, as depicted in fig 5C) and HR (t = 21.08, as indicated in fig 5C). The action potential waves alternately generated from two heterogeneous areas induced VF-like ECGs signals (as portrayed in Fig 6, with FD = 25%). In contrast to scenarios with FD less 20%, both HL and HR maintain the EAD island persistently throughout the simulation and causing subsequent ectopic beats. An anomaly occurs when t = 27.05, the EAD island in HR vanishes (Fig 5C). However, it was soon activated by the action potential wave from the HL. Throughout the entire simulation, these two competing foci sustain the VF activity. A movie of this simulation with voltage maps along with the ECG is provided as Movie S2. Interestingly, across all 2D simulations, the propagation of depolarization waves from the EAD island initially targets the Epi and Endo tissue before affecting the mid tissue outside of heterogeneous areas due to prolonged effective refractory period (ERP)

**Fig 6.**
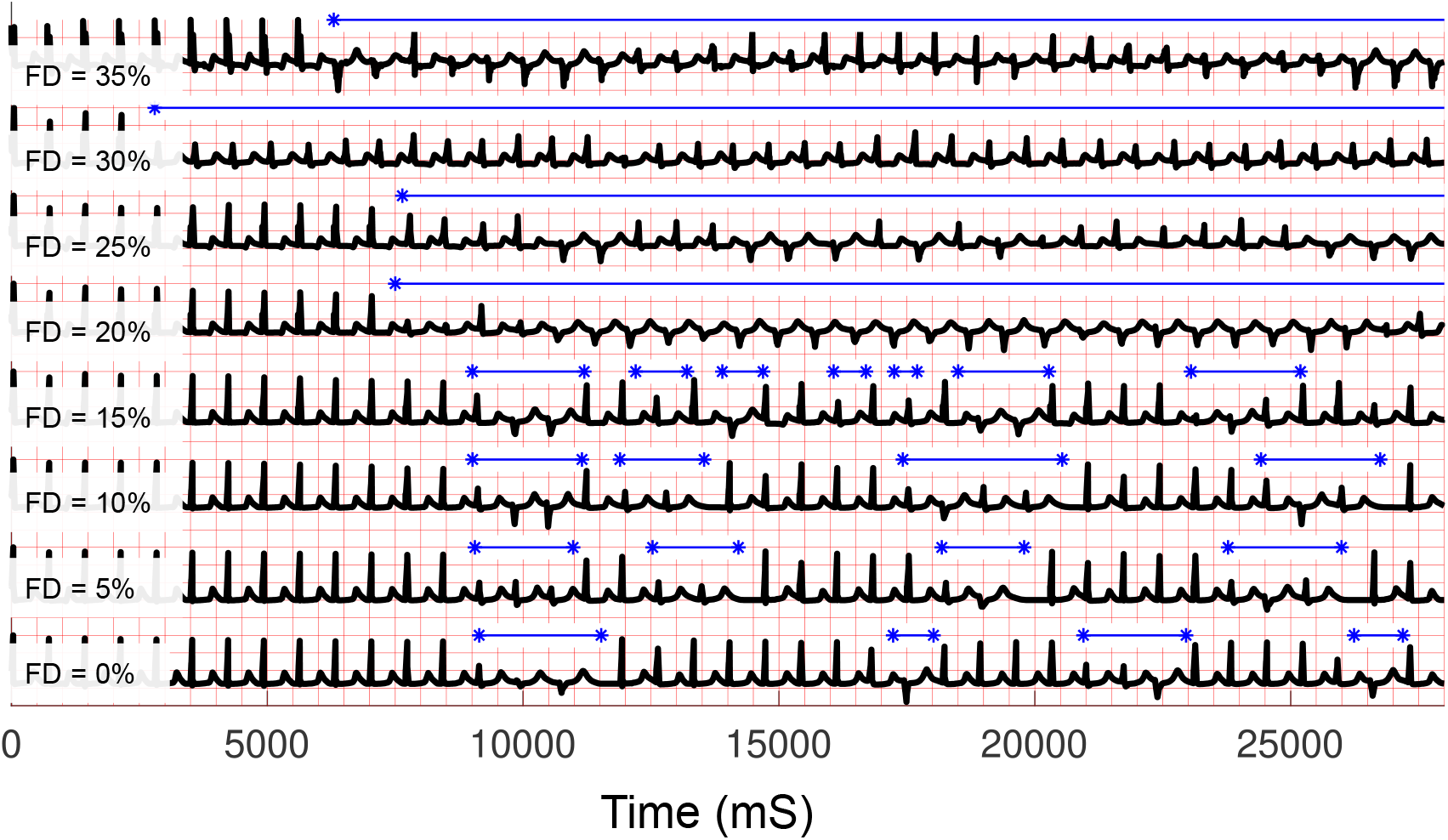
ECGs of 2D model in 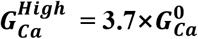 and 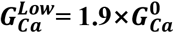, FD from 0% - 35%. Each red box length is 500ms on the x-axis. The black line is the Pseudo-ECG generated from the simulation. The amplitude of the ECG signal is normalized for ease of comparison. The blue star line is intervals of the abnormal excitations.

Fig 6 shows ECG patterns under various FD levels. The movie files of voltage map are in the Appendix S7. When the FD is under 15%, the PVCs are generated but cannot be sustained, and VF did not occur. Ectopic beats occurred intermittently, and the frequency of occurrence is not significantly related to FD. After several ectopic beats from the HL or the HR, the abnormal excitation will stop temporarily as EAD islands disappear. Then the ectopic beats begin again after several normal beats. As FD increases to more than 20%, we observed VF-like activity that lasts longer after the first focal beat occurred. The HL and HR alternately or simultaneously generate the ectopic beats from the surrounding tissue, which connects to the edge of the EAD islands. The orientation of action potential waves gradually changes, showing complex dynamics. The occurrence of PVCs and the duration of VT/VF increase with the increase of FD.

### Effects of RR

In addition to fibroblasts, RR is also a critical factor for EADs and PVC formation. Reduced RR leads to substrates for VF. In the simulations in Fig 5, the RR was set at 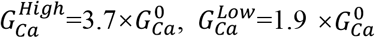 (Fig 1). Here we varied RR by changing the maximum conductance of the L-type calcium channel (G_Ca_). To reduce (or increase) RR, we adopted the approaches proposed by Panfilov, et al. (2016) [35]. We reduced the conductance of depolarizing currents *I*_*Kr*_ and *I*_*Ks*_, with the decreased values consistent with previous experiments. We also increased the conductance of *I*_*CaL*_ which responsible for EAD formation during the plateau phase of the action potential. The original value of conductance of *I*_*CaL*_ keeps consistent with the TP06 model [24], which is denoted by 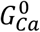. Conductance of *I*_*CaL*_ 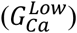 was varied from 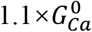 to 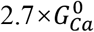, while the conductance of *I*_*CaL*_ in heterogeneous areas 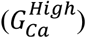 was varied from 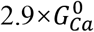 to 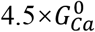.

Fig 7 shows the summary of the results. We set RR at five levels by varying G_CaL_. FD was also set into 8 levels, from 0% to 35%. We paced 40 cycles (28s at the frequency of 1.4 Hz). We recorded the proportion of abnormal excitation (fig 7A) and excitation patterns of tissue (fig 7B) in 28 s for each simulation. The specific ECG was obtained in the model shows in S8 Appendix Fig 1. When RR is high 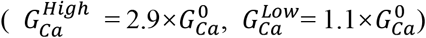, tissue cannot generate PVCs at low FD and only generates PVCs and sustained VF at high FD. Under reduced RR conditions, the total number of abnormal excitation activities increases, and duration of VF also increases. The PVCs and sustained VF occur more often until FD becomes around 30% regardless of the level of RR.

**Fig 7.**
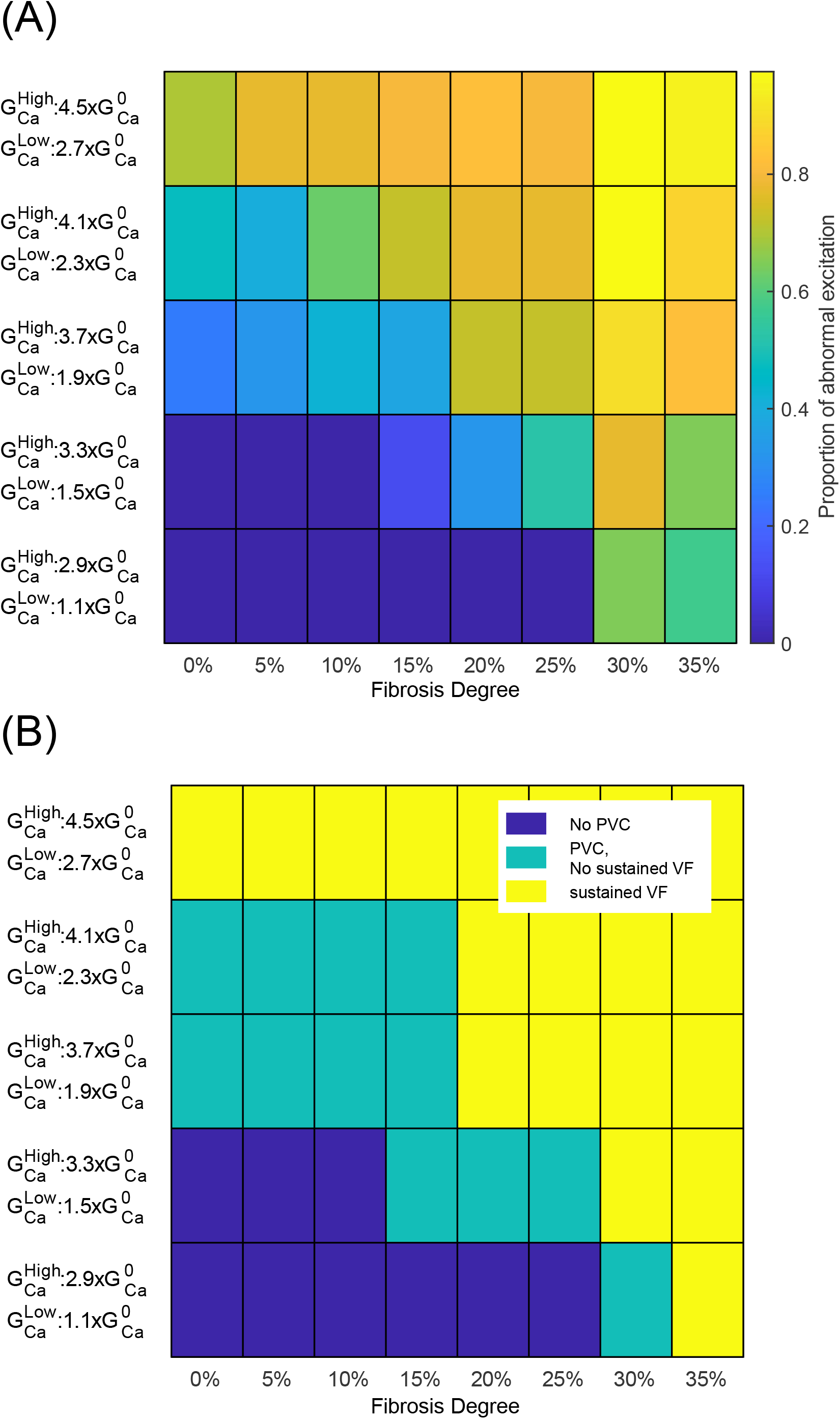
(a) Colormap of the proportion of abnormal excitation, (b) Phase map of the levels of RR and FD vs. initiation and sustain of VF. (a) The colormap shows the dependence of the sensitivity of abnormal excitation at the different levels of RR and FD. (b) The colormap shows three different dynamical states: no PVCs, have PVCs but no sustained VF, sustained VF. In the y-axis, the RR of tissue increases through the increment of PCa. In x-axis, the FD increases from 0% to 35%. In order to simulate PVC’s resetting effect on the sinus node excitation phase, the external pacing stimulus stops if there are ectopic beats and resumes if the PVC ends.

**Fig 8.**
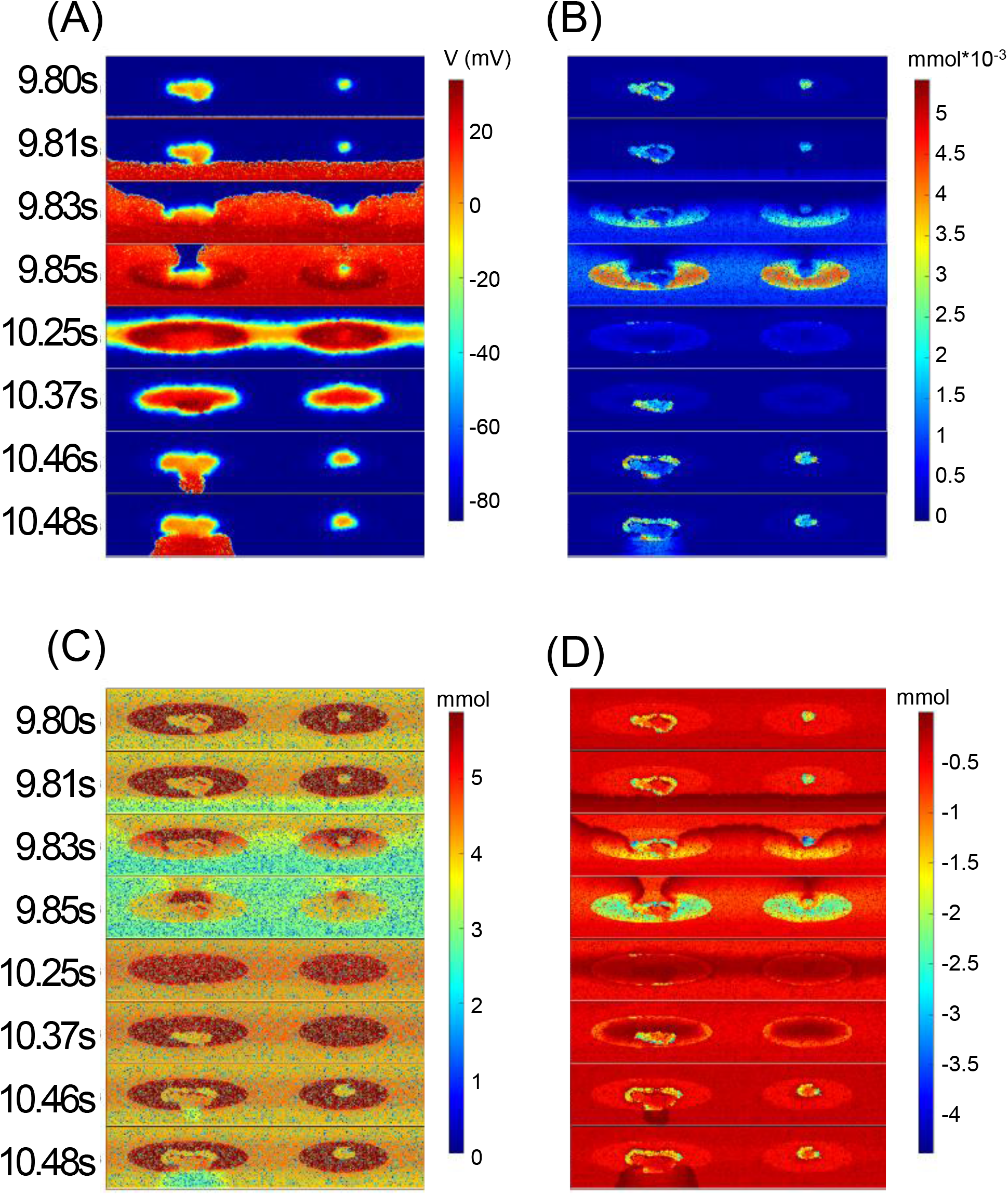
Colormap of (a) action potential (b)Cai (c)CaSR (d)INaCa from 9800ms to 10480ms.

**Fig 9.**
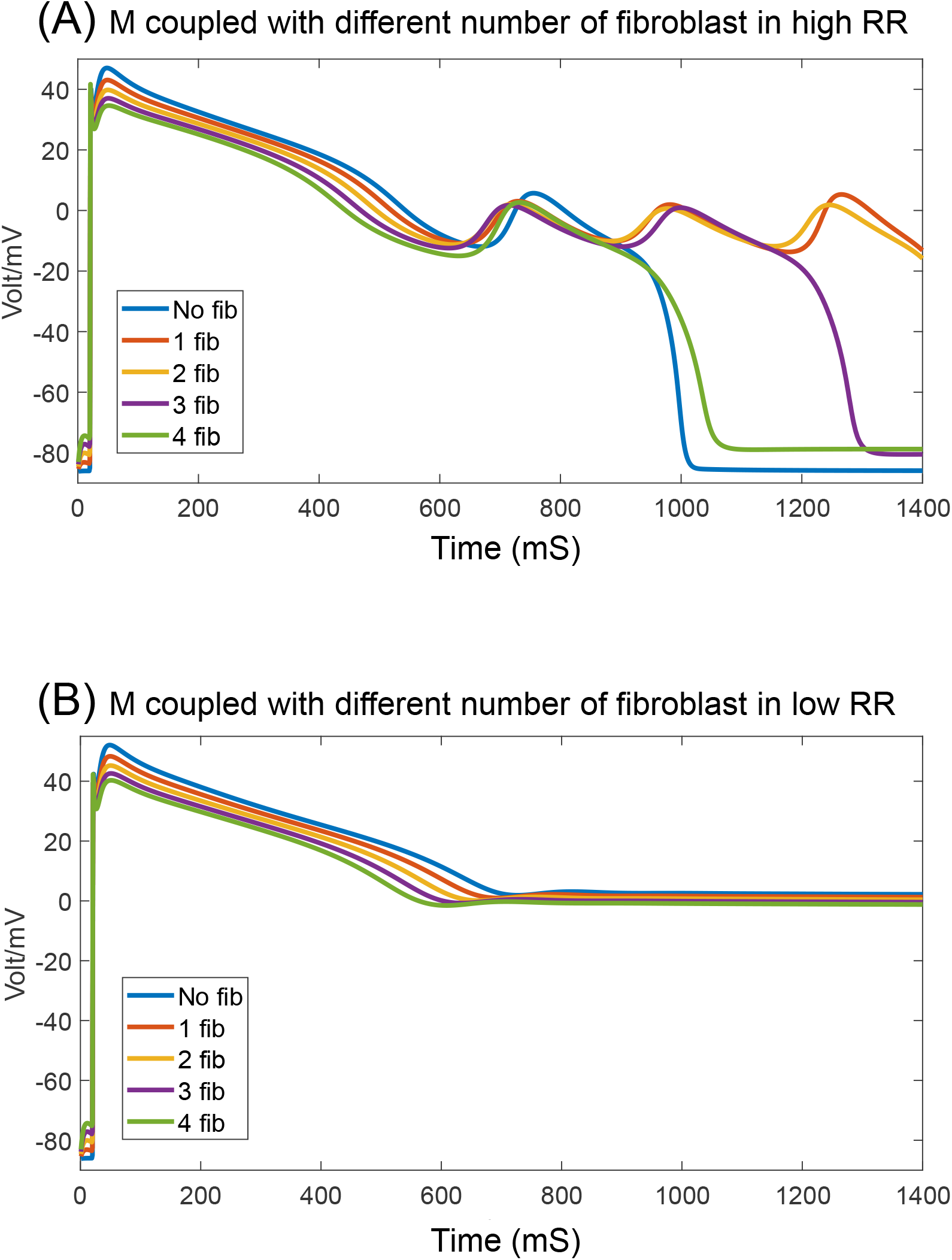
0D simulation of EAD cell’s APD in different levels of RR and different number of coupled fibroblasts.

### 3D human ventricular simulation

Subsequently, we conducted 3D simulations to investigate the initiation and sustenance of VF in a 3D cardiac model and compared the results with those obtained from 2D simulations. Here we first show the comparison between FD=15% (figs. 11A and B) and FD=35% (figs. 11C and D). In these simulations the RR is set at 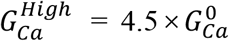 and 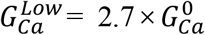 (fig 3). The total simulation time is 40 pacing cycles (= 28,000 ms) at a frequency of 1.4 Hz. The external pacing stimulus stops if there are ectopic beats and resumes if the PVC ends as same as 2D simulation.

**Fig 10.**
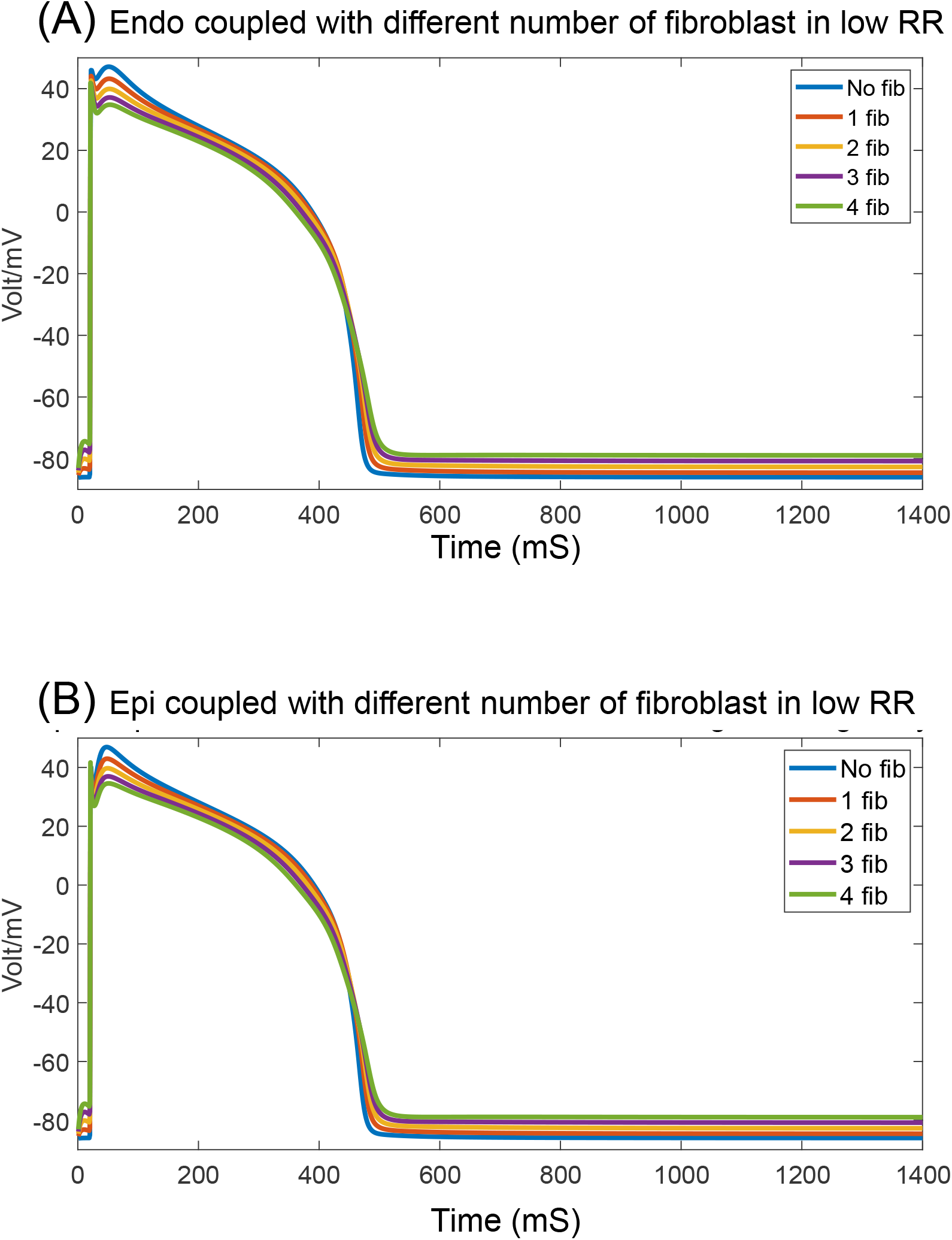
APD of (a)Endo and Epi cells with different number of coupled fibroblasts.

**Fig 11.**
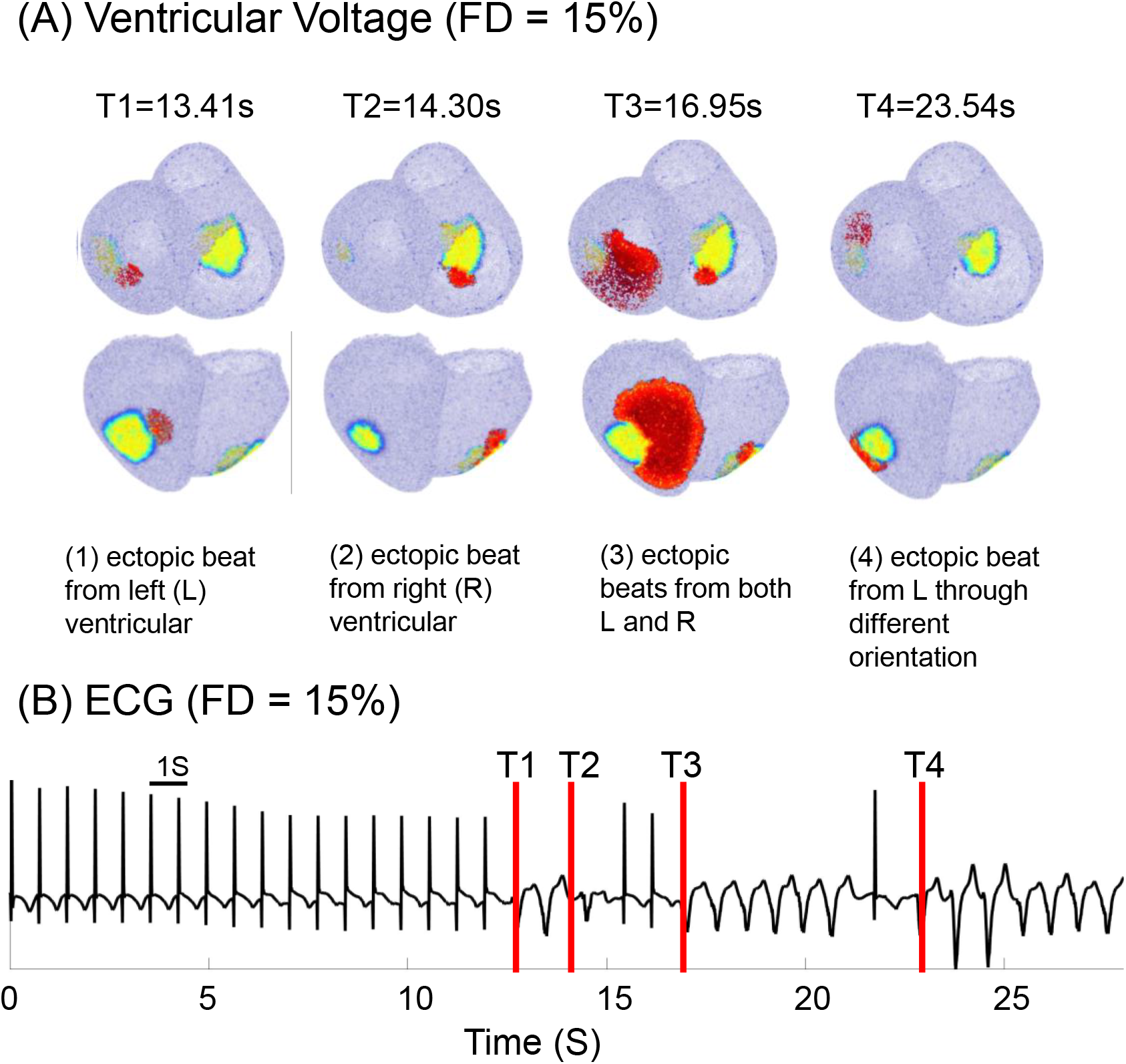
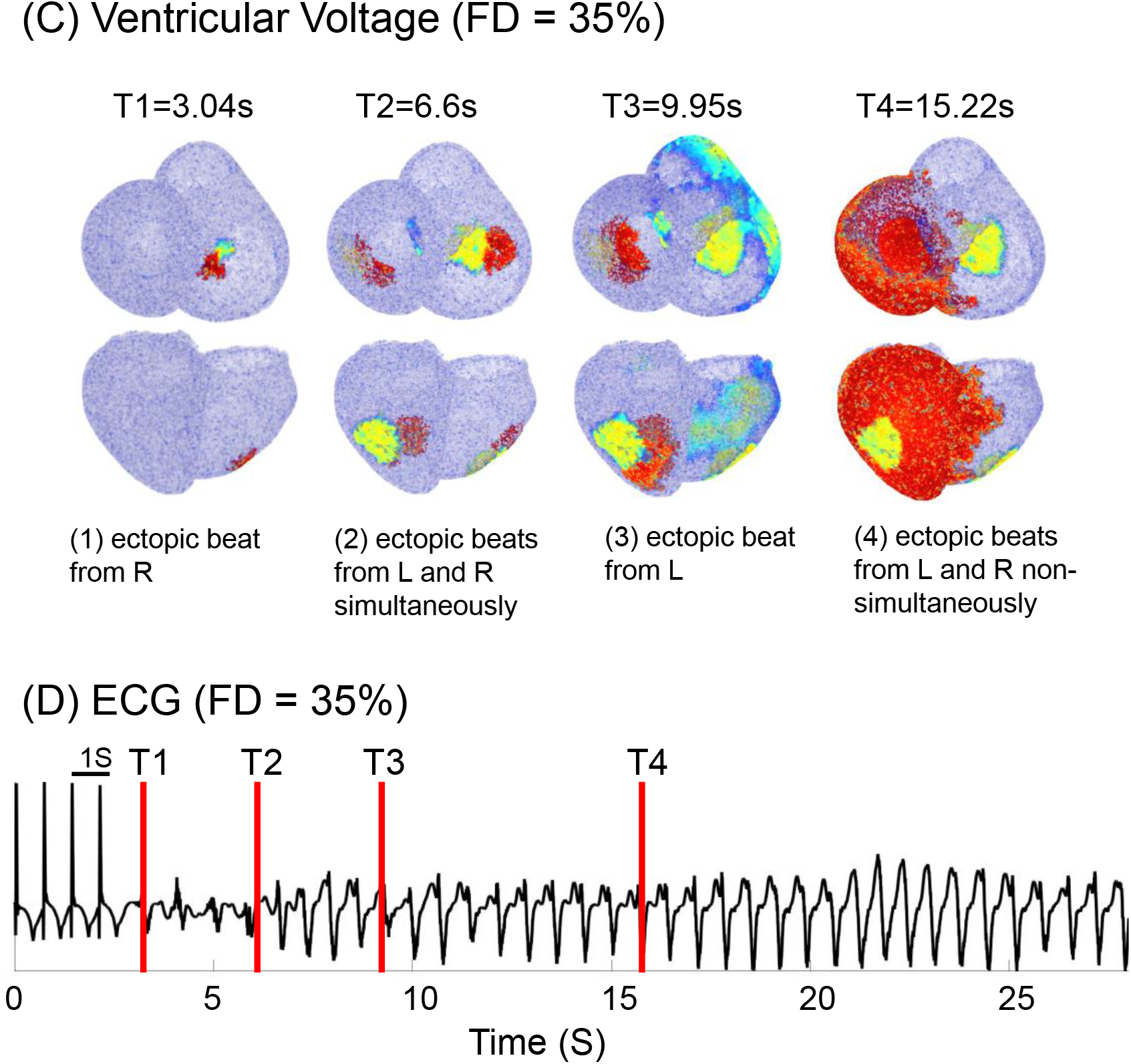
Voltage maps and ECG of 3D model in 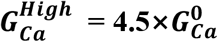 and 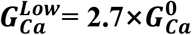, FD = 15%, 35%. (A) Voltage maps (FD = 15%), (B) ECG (FD = 15%), (C) Voltage maps (FD = 35%), (D) ECG (FD = 35%)

When FD is 15%, the first ectopic beat originates at t = 12.62 s from LV (heterogeneous area in the left ventricle, see Fig 3), Subsequently, the following ectopic beat was generated at t=13.41 s from the LV. At t = 14.30 s another ectopic beat arose from the RV (heterogeneous area in right ventricle). Then at t = 15 s the PVC terminated as two EAD islands disappeared consecutively (fig 11A). Then, after two normal pacing beats, two ectopic beats from LV and RV respectively (t = 16.95 s, fig 11A), and maintained the EAD island in the LV, abnormal excitation patterns ended at t = 21.63 s. Despite originating from the same heterogeneous area, the orientations of the depolarization waves may differ, in fig 11, the orientations direction is clockwise when t = 13.41 s but goes reverse in t = 23.54s. This variation in depolarization wave orientations contributes to changes in the amplitude of the ECG waveform, resembling a VF-like pattern (fig 11B). Similar to the 2D simulation with low FD, the abnormal excitation pattern cannot be sustained as EAD islands in both the LV and RV disappear nearly simultaneously. Consequently, the tissue intermittently returns to the normal pacing cycle. A movie of this simulation with voltage maps along with ECG is provided as Movie S3.

When FD set to 35%, the initiation of ectopic beat arises from RV (t = 3.04 s, fig 11C). After several subsequent ectopic beats generated from LV, the ectopic beats from both LV and RV simultaneously (t = 6.6 s, fig 11C). Additionally, there are instances where ectopic beats are generated non-simultaneously from the LV and RV, (t = 23.64 s, fig 11C). This complex changing excitation pattern generates a VF-like waveform (fig 12). The LV and RV can alternately or simultaneously generate depolarization waves. In essence, they become two competing foci with distinct rates in the LV and RV, akin to the experimental results demonstrated by D’Alnoncourt et al [36]. These two local heterogeneous regions sustain longer EAD and trigger PVCs successively, preventing the tissue from returning to the resting state due to simultaneous repolarization. Consequently, VF becomes self-sustaining after the occurrence of the first ectopic beat. The corresponding voltage maps for Figure 8 are provided in supplementary S3 Movie, along with the ECG.

**Fig 12.**
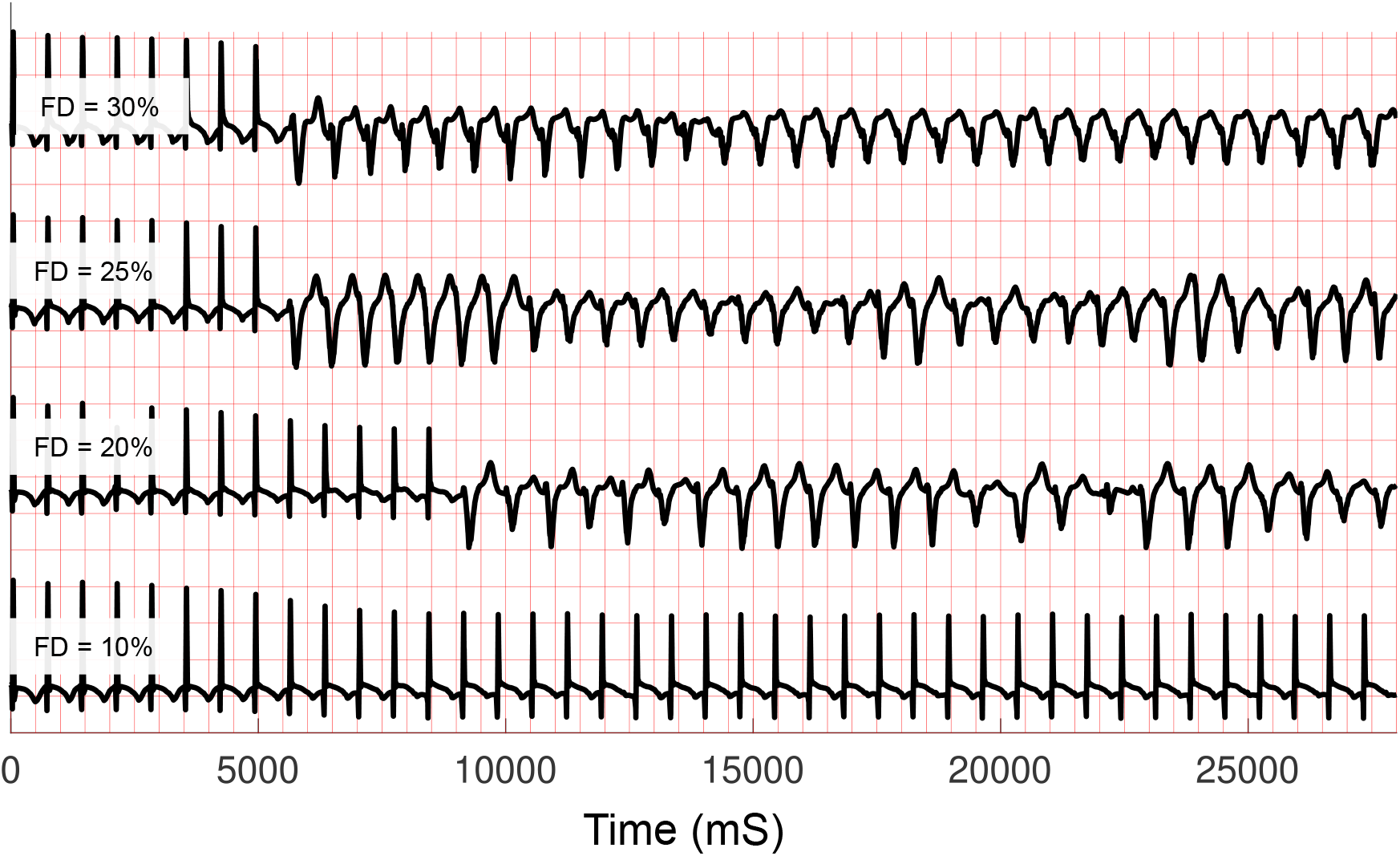
ECGs of 3D model in 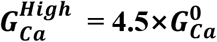 and 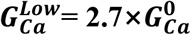, FD = 10%, 20 - 30%. Each red box length is 500ms on the x-axis. The black line is the Pseudo-ECG generated from the simulation. The amplitude of the ECG signal is normalized for ease of comparison.

In addition, these two cases, we also simulated the 3D heart model under different FD. Fig 12 shows the ECGs under various FD. When FD is set under 15%, no ectopic beat occurs. As shown in Fig 12 (FD = 10%), the tissue keeps the normal pacing cycle during the whole simulation. Although the EAD islands have been generated, no surrounding tissue can be activated before the subsequent normal pacing. When FD is 15%, the PVCs occur, but the VF-like waveform cannot maintain. When FD increases above 20%, the time point of the first ectopic beat, which initiates the VF, becomes earlier with the increase of FD. The self-sustainable VF occurs after the first ectopic beat. As we have shown in the case of FD = 35%, the abnormal excitation patterns are sustained by two competing foci. These results indicate that the PVCs and long-lasting VF occur in 3D tissue under the similar FD and RR levels as shown in the 2D simulations.

## Discussion

### Fibroblasts and EAD formation at the tissue level

In this study, we investigated the effects of fibroblasts on the formation of EADs, PVCs, and VT/VF. Firstly, the study reveals a direct relationship between FD and APD prolongation. As FD increases, APD also lengthens especially under the RR conditions, indicating a greater propensity for EADs, which are associated with ventricular arrhythmias. In the absence of fibroblasts, the APDs in the tissue were already prolonged due to the RR condition (Fig 4, 0% case). There is also variability in APD since the tissue is heterogeneous. When fibroblasts were coupled to myocytes, APD was further increased. This is consistent with findings from previous studies [16,37, 38]. When the AP was prolonged enough with the 30% of fibroblasts, EADs were also observed. Interestingly, coupling a myocyte with a small number of fibroblasts under high RR conditions promoted EAD formation, while coupling with a large number of fibroblasts facilitated the termination of EADs. These observations underscore the importance of fibrosis in modulating electrical properties within cardiac tissue, potentially leading to arrhythmogenic substrate formation. In addition to the duration, variability in APD also increased as FD increased. This is also potentially arrhythmogenic since it may lead to large gradients of the refractoriness. As discussed in the next section, the heterogeneous distribution of refractoriness and the formation of EADs due to fibroblasts play crucial roles in the formation of PVCs and thus the initiation of VT/VF.

### The roles of fibroblasts in PVC formation and maintenance of arrhythmias

Although EADs can form PVCs, EADs alone cannot form PVCs. In addition to afterdepolarizations, a gradient of refractoriness is required for the propagation of the triggered action potential.

Fibroblasts in tissue promote EAD formation. In addition, fibroblasts in tissue also lead to gradients of refractoriness, which promote PVC formation due to EADs. There are two reasons. One is that fibroblasts increase variability in APD. The other is that fibroblasts affect M-cells much more than Epi and Endo cells, especially under the RR conditions. These effects of fibroblasts lead to large gradients of the refractoriness in tissue and promote PVC formation. In Figs 5A and 5C, after all cells outside the heterogeneous regions return to their resting state, PVCs started around the boundary of these regions. Moreover, the larger size of the EAD regions results in a longer duration of AP, thus providing more opportunity for EAD M-cells to interact with endocardial cells that have just passed their absolute refractory period, thereby generating the PVC. This observation is consistent with the findings in [41,42].

However, once FD becomes too high, the generation of PVCs is reduced. Primarily, increased fibrosis results in a relative decrease in the number of myocytes. Consequently, this leads to source-sink mismatch and reduces PVC formation. On the other hand, when FD is too low PVCs will not occur because no EADs will be formed. Thus, diffuse fibrosis has biphasic effects on PVC formation.

For the maintenance of VT/VF, as FD levels increased, short-lived VT-like activity became sustained VF-like events. Recent research demonstrates that when fibroblasts are locally distributed in the tissue, the spiral wave in tissue can be attracted and possibly to dynamical anchoring around the fibrotic area [41]. In our simulations, the reentry-type VF was also observed, and the anchoring effect of the fibrotic area prolonged the duration of VT/VF.

The 3D human ventricular simulations validated the findings from the 2D models. At low FD (<15%), no ectopic beats occurred despite the presence of EAD islands. Moderate FD (15%) led to non-sustained PVCs, while higher FD (>20%) resulted in self-sustaining VT/VF.

### Excitability of fibrosis

Although experiments have shown an electrical coupling between fibroblasts and myocytes in vitro [39], no reliable results indicate electrical coupling in vivo [27]. In a recent study by Zimik *et al*. [20], demonstrates that regardless of fibroblast excitability, fibroblasts can act as a sink or source in tissue. Moreover, human iPS cell experiments in vitro show that reducing tissue RR through I_Kr_ channel blockers and increasing tissue heterogeneity by inserting non-excitable obstacles in myocytes are critical factors in reproducing VF [40]. To test whether the excitability of fibroblasts is essential for our proposed mechanisms, we also used non-excitable fibroblasts in 2D simulation, and the fibroblast arrangement is the same as the excitable one under the same FD. We ran the 2D Simulation 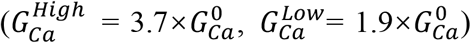 with various FD levels at 1.4 Hz for 40 pacing cycles (28 s). The results (S8 Appendix, Fig 7) showed a qualitatively similar phenomenon to the result for excitable fibroblasts. When FD is low, PVCs occur but no sustaining VF is generated. When FD increases over 20%, non-terminating VF occurs. Therefore, regardless of the excitability of fibroblasts, fibroblasts significantly affect the formation and maintenance of arrhythmias in tissue.

### Limitations

This study focused on the case of diffuse fibrosis, while there are various other fibrosis architectures, such as interstitial, patchy, and compact fibrosis. We also only studied the situation where diffuse fibrosis is randomly and evenly distributed throughout the whole tissue. However, the distribution of fibrosis in the heart is inherently uneven. When fibrosis is unevenly distributed in tissue, the abnormal excitation patterns would differ, as inferred from the results of this study. For example, if FD in the RV is too low and cannot initiate PVCs, PVC activities may occur only in the LV. We also considered only two heterogeneous areas. We can consider multiple heterogeneous areas. Since there are various (theoretically infinite) possibilities of heterogeneity, we cannot cover all of them. However, the effects of fibroblasts shown in this study would be applicable to many cases of drug-induced heterogeneity.

In this study, we considered only EADs due to reactivation of the L-type calcium channel. However, it is known that spontaneous Ca^2+^ release from the SR can also lead to EADs. DADs typically arise from this spontaneous SR Ca^2+^ release. PVCs due to these EADs/DADs are also typically observed in tissue, and fibroblasts are known to play a significant role in their development. We also did not take mechano-electric feedback into account in this study. The passive property of the fibroblast machinery affects the mechanical activity of tissues. Recently, a mechano-electric myocyte-fibroblast coupling model in-silico has been developed [43,44]. The myocyte-fibroblast simulation results show that mechano-electric feedback (MEF) is enhanced as the number of coupled fibroblasts increases. It would be interesting to investigate the relationship between MEF effects and VF at the tissue level.

## Conclusions

In this study, we have shown that diffuse fibrosis plays a complex role in the initiation and maintenance of arrhythmias. We demonstrated the interplay between the diffuse fibrosis and the drug-induced heterogeneity of APD in the genesis of ventricular arrhythmias. While a certain level of fibrosis increases the susceptibility to PVCs, excessively high fibrosis levels reduce this susceptibility. The self-sustainability of arrhythmia is also positively correlated with the level of fibrosis. These findings provide comprehensive insights into the mechanisms underlying ventricular arrhythmogenesis in the presence of fibrosis and altered repolarization dynamics. By elucidating the intricate interactions between fibroblasts, APD prolongation, and arrhythmia initiation, this study contributes to our understanding of cardiac electrophysiology and may pave the way for the development of targeted antiarrhythmic therapies. Finally, as stated in the limitations, various possibilities of heterogeneity exist. Incorporating patient-specific data into the modeling framework holds immense potential for personalized risk assessment and the development of more effective preventive and therapeutic measures for arrhythmias.

## Acknowledgments

This work was partially supported by the Japan Society for the Promotion of Science under its Grants-in-Aid for Scientific Research (KAKENHI) (no. 23H01374, 23H01382 and 22K18777) and JST Mirai JST-Mirai Program (no. JPMJMI21I1) and the National Institutes of Health

Grant R01-HL149349 (D.S.). We thank Baichuan Jiang for writing assistant of this article and Zhou Zikai for development of calculation model.

## Author Contributions

Conceptualization: Kayo Hirose, Daisuke Sato.

Data curation: Daisuke Sato

Formal analysis: Daisuke Sato

Investigation: Kayo Hirose, Daisuke Sato.

Methodology: Umezu Shinjiro, Daisuke Sato

Resources: Kayo Hirose, Umezu Shinjiro

Software: Baichuan Jiang, Daisuke Sato

Supervision: Kayo Hirose, Umezu Shinjiro, Daisuke Sato.

Writing – original draft: Kayo Hirose, Daisuke Sato.

Writing – review & editing: Kayo Hirose, Umezu Shinjiro, Daisuke Sato.

## Data Availability Statement

All relevant data are within the paper and its Supporting Information files.

## Running Head

The effect of diffuse fibrosis on Arrhythmias under drug-induced heterogeneity

## Highlights

Tissue with higher FD is more vulnerable to initiate VF under drug-induced heterogeneity. VF’s self-sustainability is also positively correlated with FD under drug-induced heterogeneity.

## Footnotes

1 *APD90 is the time it takes for the cell’s membrane potential to repolarize by 90% after reaching its peak during an action potential. It reflects the time it takes for a cell to repolarize and return to an excitable state again.

2 *APD90-APD30 quantifies the time difference between the durations of the middle phase of repolarization (30% to 90% repolarization). The cells at this stage are above the threshold for stimulating electrical excitation of surrounding cells, reflecting the potential propagation ability of the cell’s electrical excitation.

