## Supplemental Figure 5 for "Fibroblast Density is a Risk Factor for Drug-induced Arrhythmias"

### 1 Supplementary material

#### 1.1 Activation time of Purkinje network

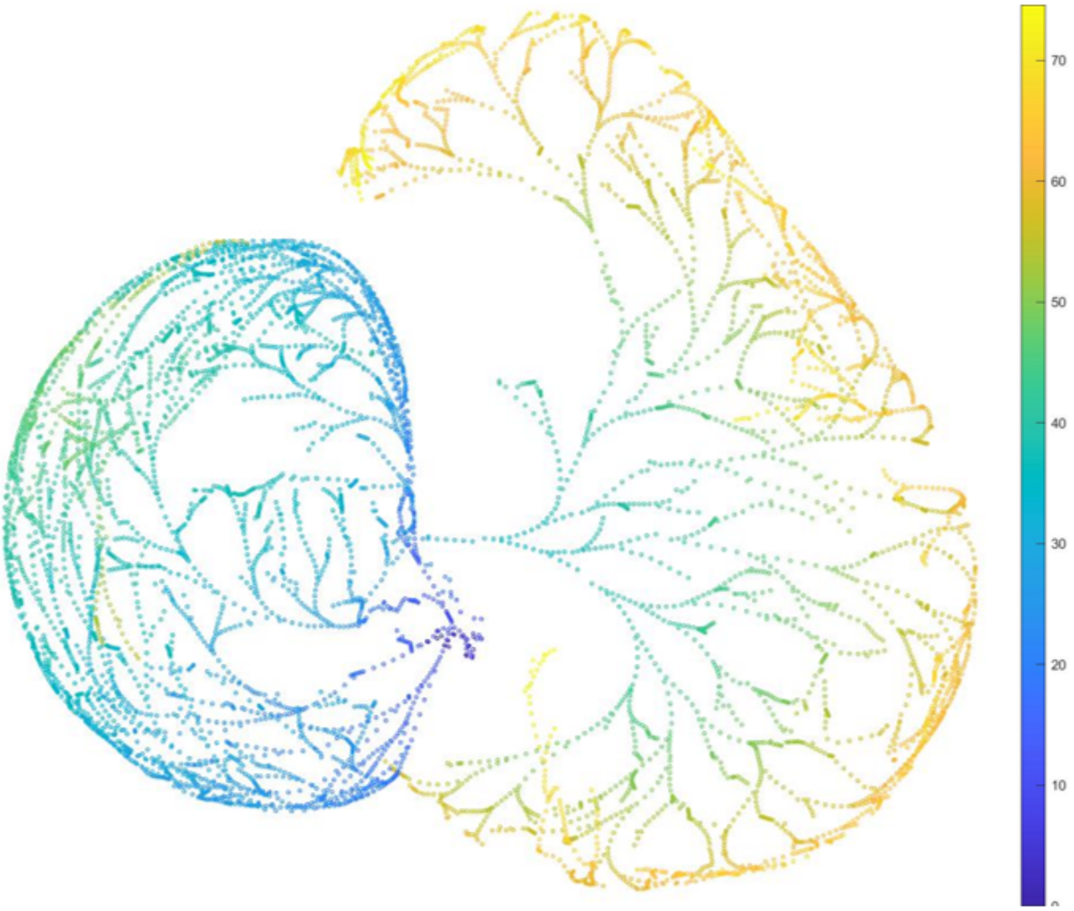


Fig 1: Activation time map for Purkinje network in the 3D ventricular model

The earliest activation time is 0.49 ms. The last activation time is 75.6 ms.

#### 1.2 2D simulation, excitable fibroblast, pacing frequency:1.4 Hz

##### 1.2.1 Levels of heterogeneity: $G_{Ca}^{High} = 2.9\times G_{Ca}^{0}$ and $G_{Ca}^{Low} = 1.1\times G_{Ca}^{0}$

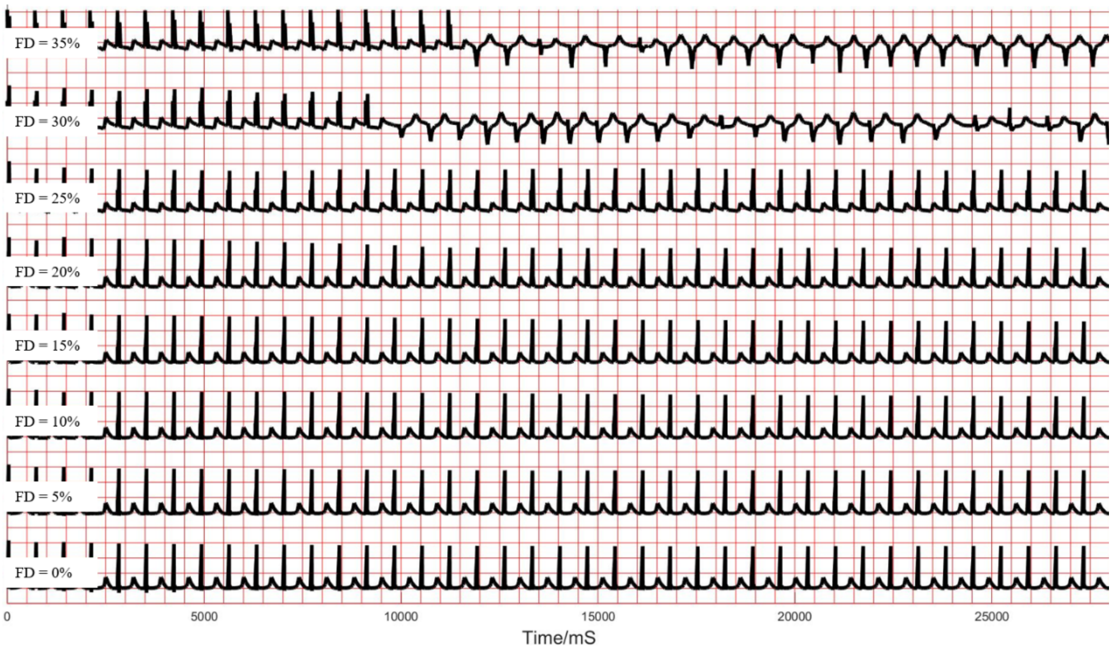


Fig 2: ECGs of 2D model in $G_{Ca}^{High} = 2.9\times G_{Ca}^{0}$ and $G_{Ca}^{Low} = 1.1\times G_{Ca}^{0}$, FD from 0% - 35%

##### 1.2.2 Levels of heterogeneity: $G_{Ca}^{High} = 3.3\times G_{Ca}^{0}$ and $G_{Ca}^{Low} = 1.5\times G_{Ca}^{0}$

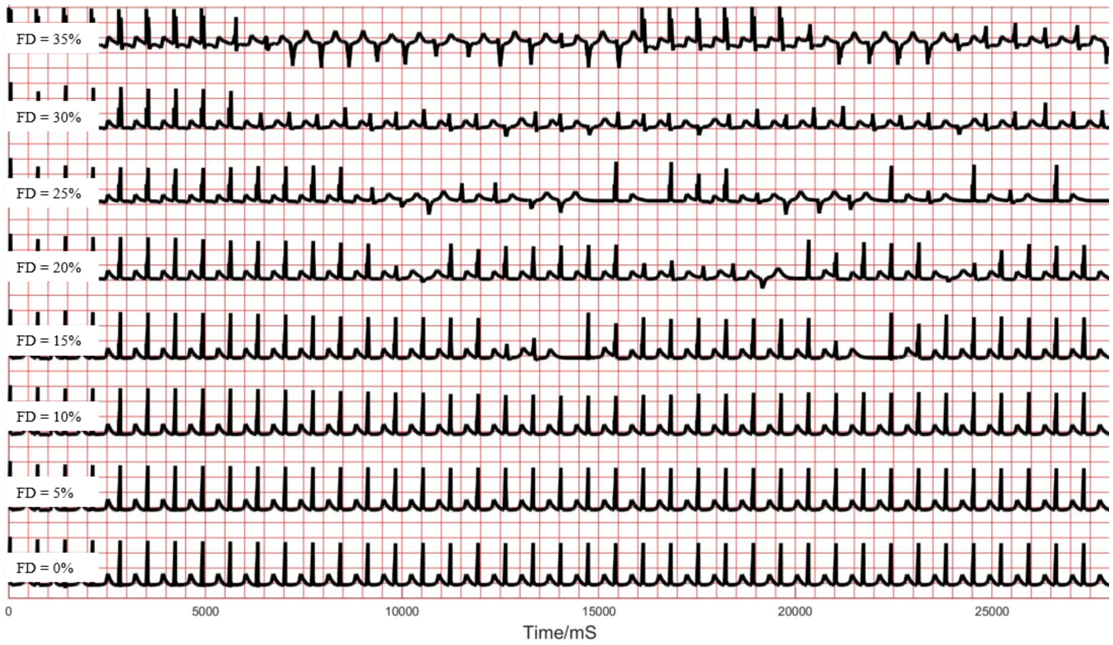


Fig 3: ECGs of 2D model in $G_{Ca}^{High} = 3.3\times G_{Ca}^{0}$ and $G_{Ca}^{Low} = 1.5\times G_{Ca}^{0}$, FD from 0% - 35%

##### 1.2.3 Levels of heterogeneity: $G_{Ca}^{High} = 4.1 \times G_{Ca}^{0}$ and $G_{Ca}^{Low} = 2.3 \times G_{Ca}^{0}$

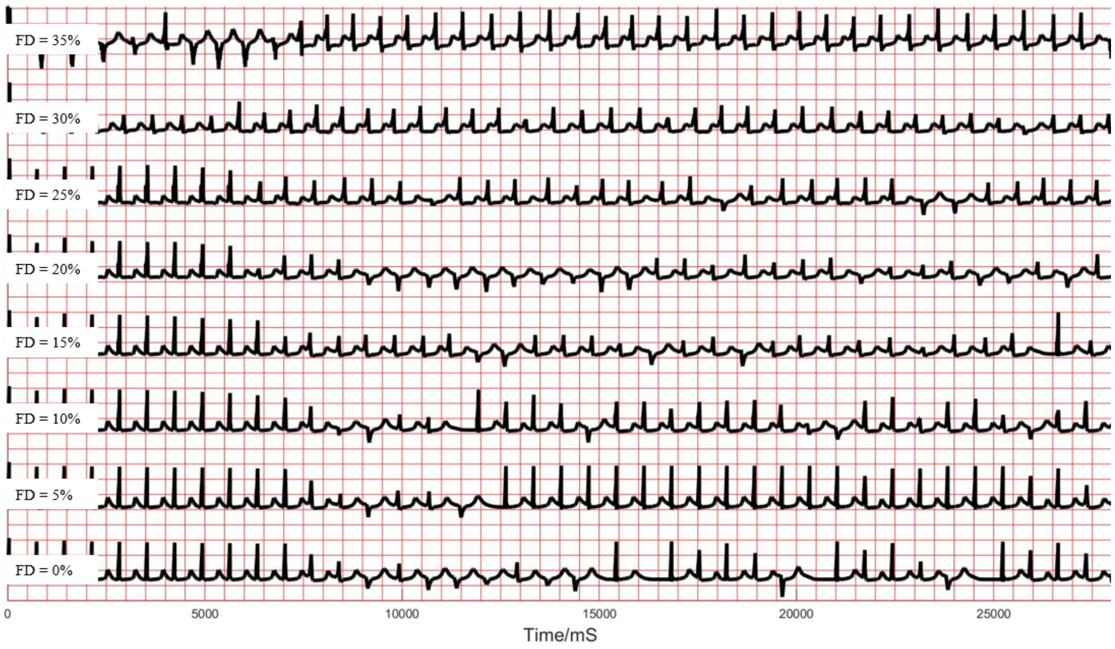


Fig 4: ECGs of 2D model in $G_{Ca}^{High} = 4.1 \times G_{Ca}^{0}$ and $G_{Ca}^{Low} = 2.3 \times G_{Ca}^{0}$, FD from 0% - 35%

##### 1.2.4 Levels of heterogeneity: $G_{Ca}^{High} = 4.4 \times G_{Ca}^{0}$ and $G_{Ca}^{Low} = 2.6 \times G_{Ca}^{0}$

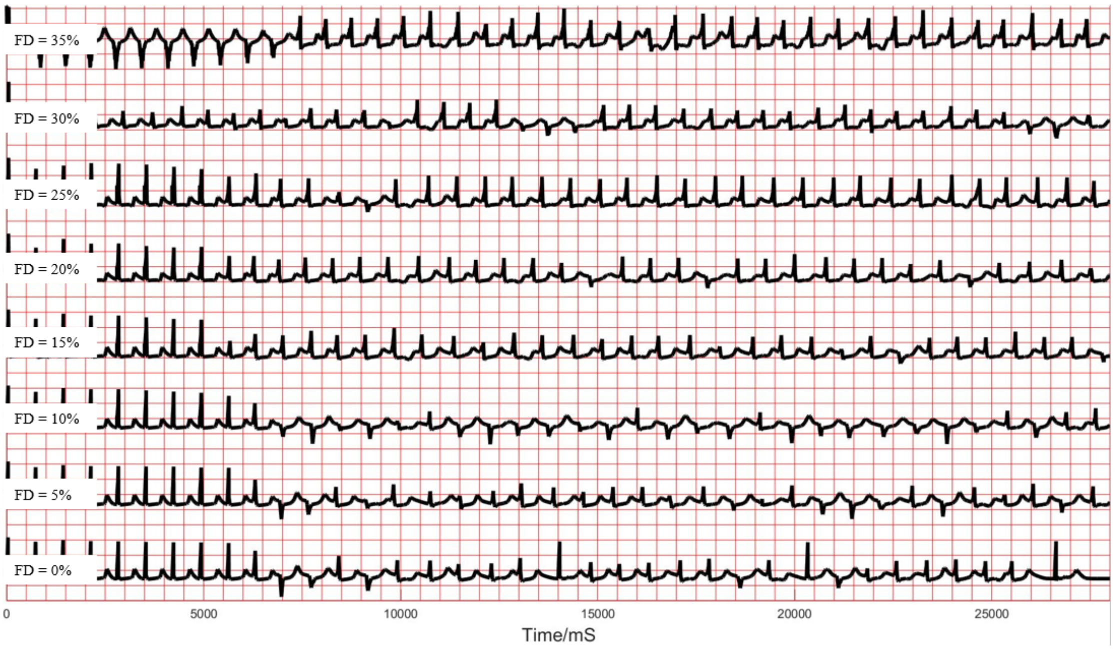


Fig 5: ECGs of 2D model in $G_{Ca}^{High} = 4.4 \times G_{Ca}^{0}$ and $G_{Ca}^{Low} = 2.6 \times G_{Ca}^{0}$, FD from 0% - 35%

**1.3 2D simulation, level of heterogeneity: $G_{Ca}^{High} = 3.7\times G_{Ca}^{0}$, $G_{Ca}^{Low} = 1.9\times G_{Ca}^{0}$, excitable fibroblast, pacing frequency: 2 Hz**

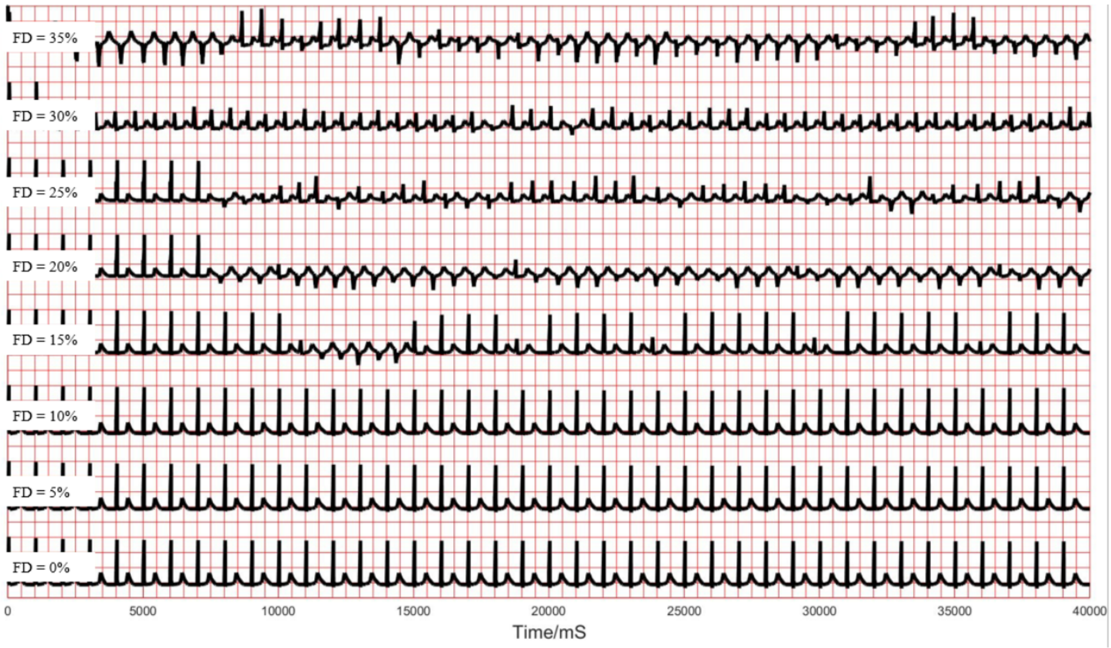


Fig 6: ECGs of 2D model, FD from 0% - 35%

**1.4 2D simulation, level of heterogeneity: $G_{Ca}^{High} = 3.7\times G_{Ca}^{0}$, $G_{Ca}^{Low} = 1.9\times G_{Ca}^{0}$, non-excitable fibroblast, pacing frequency: 1.4 Hz**

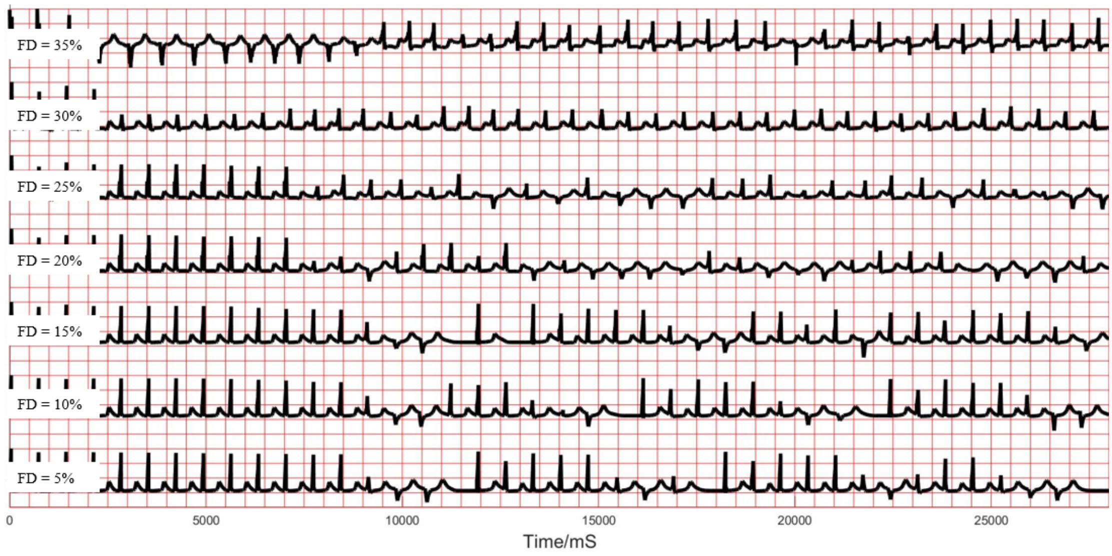


Fig 7: ECGs of 2D model, FD from 5% - 35%
